# A Functional and Non-Homuncular Representation of the Larynx in the Primary Motor Cortex of Mice, a Vocal Non-Learner

**DOI:** 10.1101/2024.02.05.579004

**Authors:** César D. M. Vargas, Rajvi K. Agravat, Elena N. Waidmann, Christodoulos Bochalis, Hector Bermudez, Theodoros Giannakopoulos, Erich D. Jarvis

**Author notes:** Equal contribution (Rajvi K. Agravat, Elena N. Waidmann).

## Abstract

Vocalization is a complex behavior ranging from fully innate to advanced vocal learning. Vocal learning species possess a vocal primary motor cortex (M1) region that makes direct projections to brainstem vocal motor neurons, which are thought to facilitate learning and fine modulation of vocalizations. Surprisingly, a similar, albeit sparse, direct projection from M1 was found in mice. Mice produce ultrasonic vocalizations (USV) which appear to be mostly innate. Modulation of these USVs is impacted by lesions to this M1 region, but genetic ablation of the cortex leads to few, if any, changes to USVs. It remained unclear whether M1 has any control over the vocal organ in a vocal non-learning species. In the current study, we found that stimulation in different parts of M1 in mice can generate contractions in laryngeal and jaw muscles, with different latencies suggestive of both direct and indirect projections to brainstem vocal motor neurons. Viral tracing reveals both single- and double-labeled populations of cortical neurons that simultaneously innervate laryngeal, jaw, and forelimb motor circuits. Chemical lesions reveal that an anterolateral orofacial region of M1 regulates the number of syllables in vocal sequences. Our results provide evidence that M1 in a vocal non-learner has some influence on vocal musculature, consistent with the continuum hypothesis of vocal learning. They also reveal that the representations of muscles for different behaviors across mouse M1 are more intermixed than previously considered. We discuss how these results impact hypotheses on the evolution of cortical vocal control and motor cortex organization.

## INTRODUCTION

The human ability to produce and finely articulate learned vocal sounds, including speech, is rare and occurs by a process of vocal learning. This behavior in songbirds and human is controlled by a network of interconnected forebrain regions, which include the human dorsal and ventral laryngeal motor cortex regions (dLMC and vLMC) located within the primary motor cortex (M1; **Figure 1A**)^1,2^, which has been argued to be involved in our human capacity to volitionally control production of learned sounds ^3–6^. The analogous circuit in vocal learning avian species appear to be convergently evolved with humans (**Figure 1B**)^2,3,7^. Humans, as well as some other animals, have evolved cortical motor neuronal cells (CMs) from layer 5 of M1 that form direct, monosynaptic projections with motor neurons in the brainstem or spinal cord. Human dLMC and vLMC have CMs (presumably from layer 5) that project to motor neurons in the nucleus ambiguus (Amb), which controls laryngeal muscles (**Figure 1A**)^8,9^. Stimulation of the vLMC^10,11^ or dLMC^12^ results in laryngeal contractions. Bats, which represent another clade of vocal learning mammals, have recently been shown to also possess these direct projections from M1^13,14^. Songbirds have analogous direct projections from the robust nucleus of the arcopallium (RA) to syringeal motor neurons in nucleus XII in the brainstem (**Figure 1B**)^15,16^. When RA is stimulated, it is possible to produce both learned and innate vocalizations^17^. Greater numbers of CMs have been hypothesized to endow a species with proportionally greater dexterity over the muscles of that circuit^18^.

**Figure 1.**
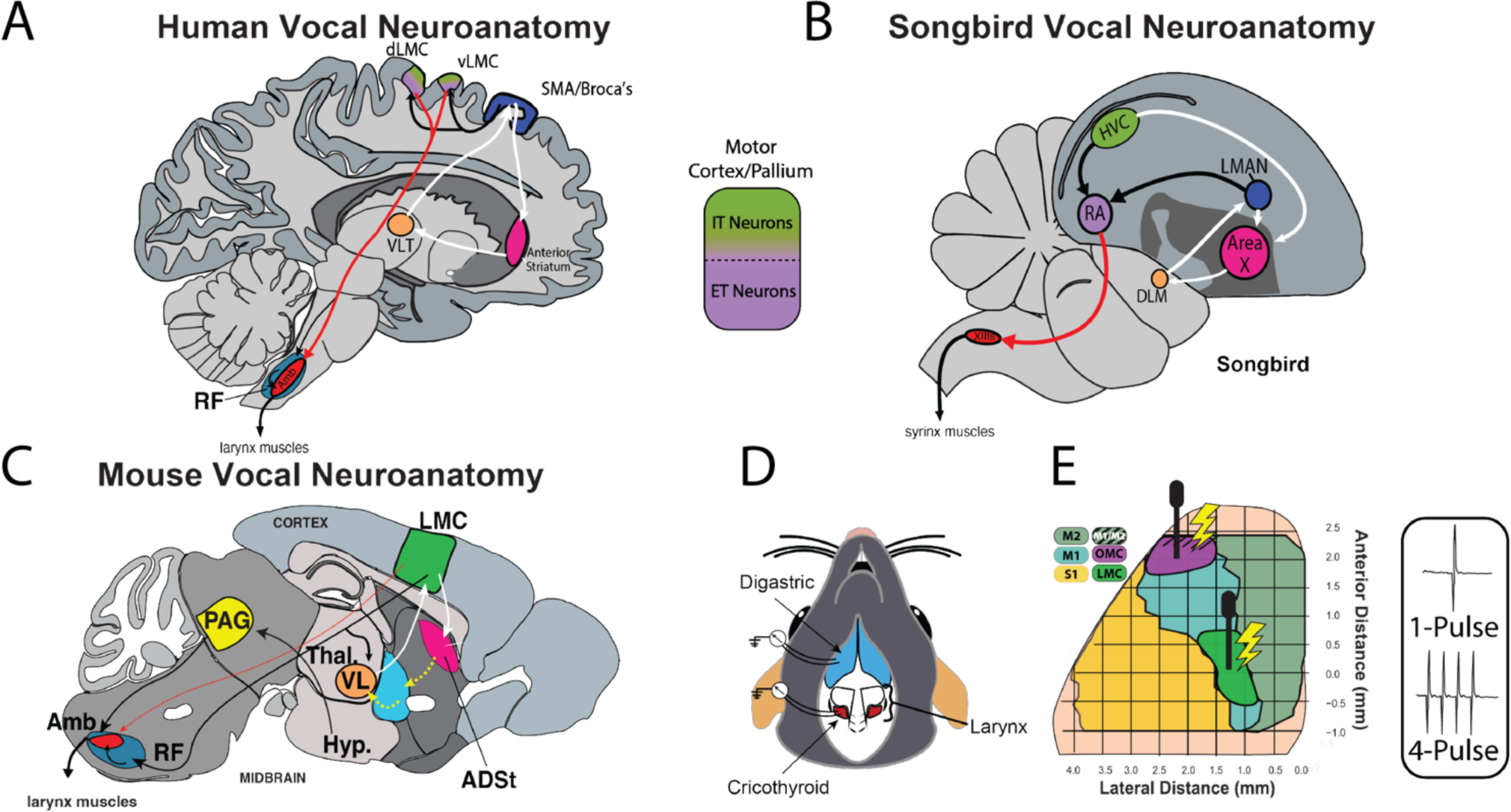
Mouse ultrasonic song system and experimental outline. (A and B) Overview of the known anatomy of the human and songbird vocal production and vocal learning system. Red arrow, direct projection from the cortical region to vocal motor neurons in the brainstem. White arrows, cortico-striatal-thalamocortical loop. Black arrows, other connections of vocal circuitry. Similar colors reflect analogous roles in the respective vocal motor and/or vocal learning circuits. (C) Overview of mouse vocal motor circuitry including sparse direct projection (thin red arrow). Yellow arrows, proposed connections for the corticostriatal-thalamic loop. (D) Overview of the two muscles from which EMGs were recorded in this study, the anterior digastric (blue) and cricothyroid (red). (E) Dorsal view of the surface of mouse cortex, with the LMC (green) and OMC (magenta) regions highlighted, and the two types of stimulation used in this study. Striped area represents where M1 is less dorsal or below M2 in the dorso-ventral axis. Anatomical abbreviations: ADst, anterior dorsal striatum; Amb, nucleus ambiguus; DLM, medial dorsolateral nucleus of the anterior thalamus; HVC (proper name); LMAN, Lateral magnocellular nucleus of the nidopallium; d/vLMC, dorsal/ventral laryngeal motor cortex; LMC, laryngeal motor cortex; M1, primary motor cortex; M2, secondary motor cortex; OMC, orofacial motor cortex; PAG, periaqueductal grey; RA, robust nucleus of the arcopallium; S1, primary somatosensory cortex; RF, reticular formation; SMA, supplementary motor area; VLT and VL, ventrolateral thalamus; XIIts, 12^th^ motor nucleus, tracheosyringeal part.

Thus far, only in vocal learners do stimulations of the primary motor region (human M1 and songbird HVC or RA) lead to vocal muscle contractions or vocal production^6,19–22^. However, stimulating the ventral premotor cortex, Area 6V, in non-human primates can induce vocal muscle movement^23–25^. Lesions in human M1 or songbird RA deteriorates or abolishes the ability to produce previously learned vocalizations, while leaving innate vocalizations intact^6,26,27^. In contrast, lesions in M1 of non-human primate species do not appear to affect vocal abilities^19^. Taken together, it had been argued that: 1) these direct-projecting cortical circuits for vocal muscles are a unique evolutionary feature of vocal learners^3,8,19,28,29^; 2) that vocal non-learners have an absent LMC and associated vocal M1 CMs; and 3) that these species mainly rely on brainstem vocal circuits to produce innate vocalizations^19,28^.

This view was more recently challenged by work in lab mice (*Mus musculus*) from our group^30^. This study found a region in M1 with vocalizing-driven immediate early gene activity that also contained a sparse direct projection to Amb (**Figure 1C**); and lesioning this region caused some degradation of frequency modulation^30^. Based on these findings, they termed this M1 region a putative, but rudimentary LMC in mice. Later work in Alston’s singing mouse (*Scotinomys teguina)* showed that they also have an LMC region that is anatomically similar to *M. musculus,* demonstrated with similar retrograde transsynaptic tracing from larynx muscles^30,31^. In a different study, in *S. teguina*, manipulations of a more anterolateral region of M1, the orofacial motor cortex (OMC), was shown to modulate vocal turn taking in a social encounter^32^. Most recently, work in two non-human primates, marmosets (*Callithrix jacchus*) and macaques (*Macaca mulatta* and *Macaca fascicularis*), also used a retrograde transsynaptic virus injected into laryngeal muscles and found labeled M1 neurons^33^. However, based on the timing of the tracer transport, they inferred that M1 only had indirect projections to Amb motor neurons. The findings in lab mice led to the continuum hypothesis of vocal learning^3,29,34^. Traditionally, vocal learning is described as an all or none trait. The continuum hypothesis proposes that rodents, non-human primates and the other so-called “vocal non-learning” species display, a subset of abilities and mechanisms present in advanced vocal learners that span a range between the dichotomous options.

Here we asked if the rudimentary LMC of M1 in mice can control vocal muscles (i.e. larynx), and if so, whether it functions primarily through a direct or indirect projections to Amb. We find that electrical stimulation of mouse LMC population can cause laryngeal muscle contraction, but at latencies predominated by indirect projections, with a minority by direct projections. In contrast, OMC has a strong influence on jaw muscles through a predominantly direct projection. Further, stimulations at some cortical locations activates both muscles. Using retrograde transsynaptic tracing shows that multiple muscles exhibit a large degree of overlapping representations in the LMC region of M1. Based on these results, we argue that cortical control of vocal muscles is more widely distributed across mammalian species than previously believed, providing further support for the continuum hypothesis of vocal learning, and that the mouse M1 has a less homuncular organization than previously believed.

## RESULTS

### Stimulation of mouse motor cortex generates vocal and jaw muscle contractions

We performed intracranial microstimulation (ICMS) by inserting stimulating electrodes in M1 while recording EMG signals from the laryngeal cricothyroid muscle (CT) and the jaw opening digastric muscle (DG) simultaneously in each animal (**Figure 1D and E and Figure S1A-C**; *n* = 7 mice). We stimulated the cortex with either 60 consecutive trials of single pulses or 30 consecutive trials of four pulses, and then calculated the average of the EMG activity, called the stimulus triggered average (StTA). This averaging helped abrogate the EMG signals produced by breathing rhythms (**Figure S1C**), and helped ensure more accurate detection of reliable EMG responses from the 60 or 30 consecutive stimulations. We manually curated StTAs plots to eliminate any with spurious electrical artifacts. Muscle activity was considered a response if the EMG activity was 2 standard deviations (2SD) above baseline before cortical stimulation for at least 0.5 ms (see Methods for more details). We note that single pulses are more rarely performed in studies of muscle EMG activity or inducing muscle movement, and even our four-pulse stimulations are far shorter (∼8.5 ms) than is typically used in other ICMS studies, which range from 40 ms to 3 s^24,25,35,36^. However, we felt it was necessary to have a conservative experimental condition for detecting direct cortical to motor neuron-muscle activity.

We stimulated the OMC region in all 7 mice, and the LMC in 4 of these. We found that single-pulse stimulation in some sites of the LMC caused contractions in the CT muscle (sites in n = 2 of 4 mice tested; example site in **Figure 2A**, individual tracers in light color, mean in bold; also see **Figure S2A**). In contrast, single-pulse stimulation in the OMC cortical region resulted in a much smaller amplitude response in the CT muscle (sites in n = 4 of 7 mice tested; example in **Figure 2B, Figure S2B**). The opposite occurred for the DG muscle, with smaller amplitude and fewer responses from LMC stimulation (sites in n = 1 of 4 mice tested) and robust responses from OMC stimulation (sites in all n = 7 of 7 mice tested; **Figure 2C and D, Figure S2C and D**). To visualize the pattern of response sensitivity within M1, we calculated a kernel density estimate weighted by the percent of times the stimulated sites were responsive across all M1 sites tested (n = 17 sites tested; **Figures S3A and B**) across all 7 mice. This visualization revealed that CT contractions occurred relatively more often from stimulations in the LMC region, whereas DG contractions occurred more often from stimulations in the OMC region (**Figure 2E and F**).

**Figure 2.**
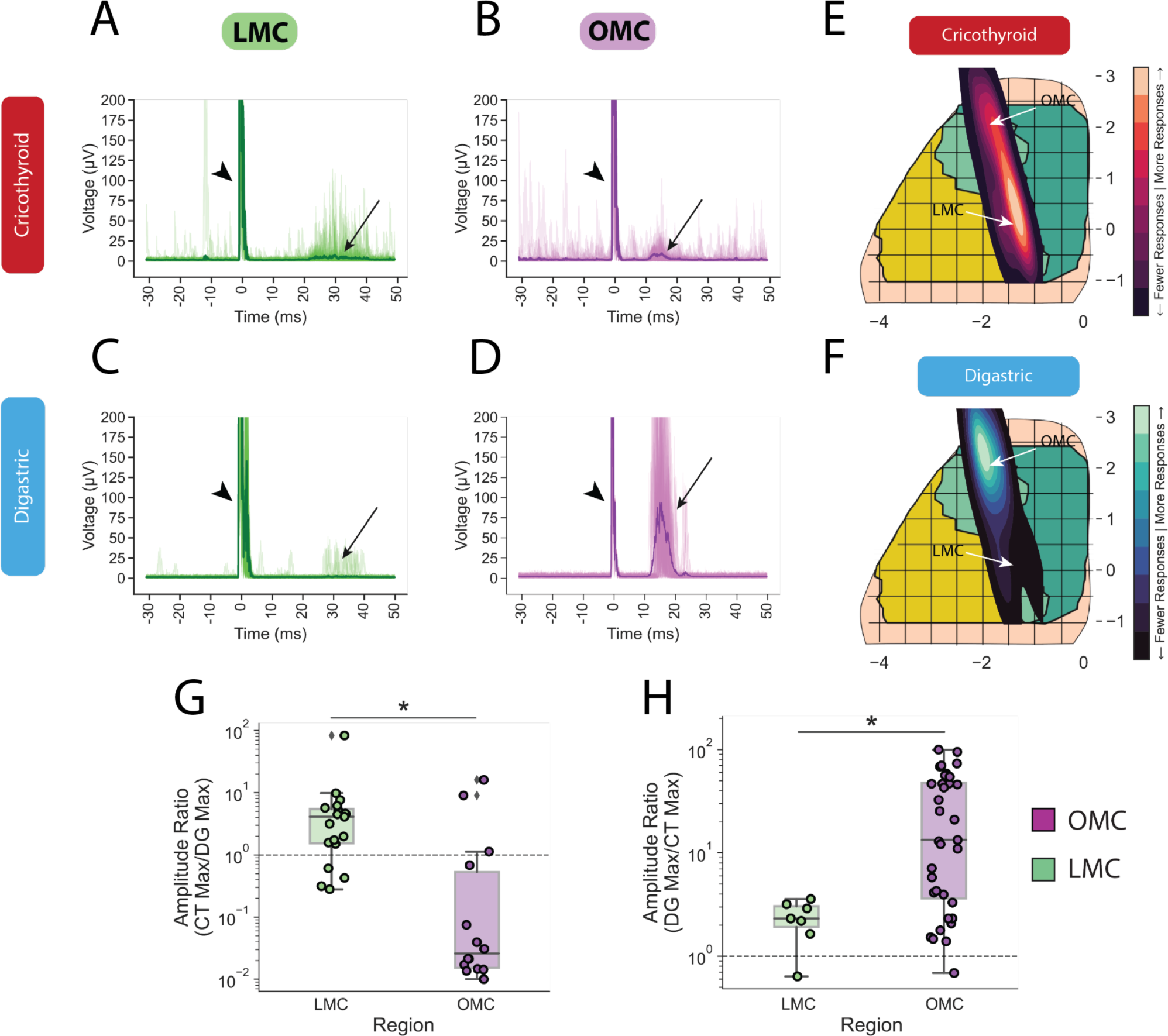
ICMS in M1 induces EMG in vocal musculature. (A and B) Example EMG responses from the CT muscle after single-pulse stimulation in the LMC (A) and OMC (B) cortical regions. Shown is a StTA plot from a single round of stimulations, at one site, in one animal. Fine traces are overlays on individual ICMS trials, bolded trace is the mean of the individual traces represented in each plot. Arrowhead, stimulus artifact. Arrows, EMG response. (C and D) Example EMG responses from the DG muscle after a single round of single-pulse stimulation in the LMC (C) and OMC (D) cortical regions. (E and F) Kernel density estimate of responsive sites weighted by percent of times the CT (G) and DG (H) were responsive at each site. (G and H) Amplitude response ratios calculated for CT/DG (E) and for DG/CT (F). CT/DG amplitude ratios were significantly higher after stimulation in LMC compared to OMC (Mann-Whitney, p=0.001). DG/CT amplitude ratios were significantly higher after stimulation in OMC compared to LMC (Mann-Whitney, p=0.027). Dashed lines denote a ratio of 1.

To quantify the differences in the response properties of the different muscles from either LMC or OMC stimulation, we calculated a response ratio, which we defined as the ratio between the maximum voltage value (µV) of the StTA when the CT was responsive (CT/DG) or when the DG was responsive (DG/CT). When stimulating the LMC region, the CT muscle’s amplitudes were on average ∼4 times higher than the DG muscle (**Figure 2G**; Mann-Whitney p=0.001). The opposite was true when stimulating the OMC region, where the DG responses that were on average ∼12 times higher than LMC stimulation (**Figure 2H**; Mann-Whitney p=0.027). We compared the minimum necessary currents we used to detect a responsive StTAs in the muscles for the coordinates within LMC and OMC, and found that there was no significant difference in the minimum current (i.e. sensitivity) needed to generate an EMG response in either muscle (**Figure S3C and D**; Mann-Whitney p>0.05).

Within the same mice, we also used four-pulse stimulations in both LMC (6 mice stimulated) and OMC (7 mice stimulated) and found similar patterns of response properties in terms of amplitude differences for each muscle (**Figure S4A-D**). Four-pulse LMC stimulation elicited consistent CT responses in 5 of 6 mice, and DG responses in 3 of 6 mice; four-pulse OMC stimulation elicited CT responses in 6 of 7 mice, and DG responses in 7 of 7 mice. We again performed a weighted kernel density estimate calculated on the responsive StTAs derived from four-pulse stimulations (n = 45 sites tested; **Figure S4E and F**), which revealed a similar, but broader pattern relative to single-pulse stimulations (**Figure S4G and H**). This included responses from S1 when tested in three of the mice. Further, some of our responsive sites overlapped with previously described representations of forelimb musculature^36^. To test this, we again performed ICMS while recording from CT, DG, and the extensor carpi radialis (ECR, a forelimb muscle) simultaneously (*n =* 1 animal). Using four-pulse stimulations in M1, we found responses in CT, DG, and ECR muscles simultaneously (**Figure S4I-K**).

To better understand the differences in the EMG responses from ICMS in different cortical regions, we performed a low dimensional reduction of all StTAs using uniform manifold approximation (UMAP). Examining the single pulse stimulations, the UMAP separated mainly by muscle identity (CT and DG; **Figure S5A**), not cortical region identity (OMC or LMC; **Figure S5B**), and this was the case whether there was a muscle response (black circle edges) or not (no edge). There was a small cluster of sites in OMC that were specific to DG responses (**Figure S5A and B, circled**). We performed K-Means clustering of the StTAs using n = 4 clusters corresponding to the four muscle-cortical region pairs of interest (CT-OMC, CT-LMC, DG-OMC, and DG-LMC) and overlaid cluster identities on the UMAP, showing these were closer to the muscle separation than the cortical separation, although still different than both (**Figure S5C**, each color is a different K-Means cluster). The four-pulse stimulation results were similar, with a subset of responses from the DG muscle appearing as more unique and dominating one of the k-means clusters (**Figure S5D-F**, circled).

Overall, our stimulations reveal a functional role of the previously identified putative LMC neurons for inducing laryngeal muscle movements^30^ and some activation of the DG. The results also support OMC’s previously described role in DG movements^32,37^ and, to a lesser extent, a role in controlling CT movements. The increased ability to identify more consistent cortical sites within a region and in more animals when using four-pulse versus single-pulse stimulations suggest a heterogeneity in connectivity and potential states.

### Synaptic latencies indicate direct and indirect connectivity

Considering heterogeneity, if the LMC projection onto Amb motor neurons is primarily monosynaptic in mice, one would expect a short ∼10 ms latency responses in the CT muscle contraction; if it were primarily indirect, the latencies would be ∼15 ms or greater^20,38,39^. We noted latencies that included these ranges from both LMC and OMC stimulation to both CT and DG muscles, with consistency in the 60 consecutive single-pulse stimulations (**Figure S2A-D**). In some cases, only a few of the 60 trials showed a response, which tended to be variable in their latency (**Figure S2C**). To quantify stimulation response latencies, we first defined response latency in positive StTAs as the point where the average signal across the 60 trials crossed our 2SD threshold criterion for at least 0.5 ms. We plotted a histogram of the distribution of latencies from responsive StTAs across all 7 animals tested with single-pulse ICMS. We found that the LMC latency to CT muscle response ranged from ∼5 to 28 ms, with a mean of 16.77 ms; whereas those from stimulation in OMC to CT muscle response ranged from ∼5 to 18 ms with a mean of 10.51 ms (**Figure 3A**). We plotted the latencies as cumulative distributions and found that stimulation in LMC results in ∼50% of latencies shorter than 15 ms to CT, while the rest were longer; OMC stimulations resulted in all but one latency measurement, 97%, being less than 15 ms to CT (**Figure 3B**). We found similar latency results from OMC stimulation and the DG responses, with a mean of 10.35 ms (**Figure 3C-D**). The one animal that showed LMC stimulation to a DG response with single pulses had a wide range of distributions in timing, with a median 16.61 ms, which is still longer than the response from OMC stimulation. That is, overall, latencies to the CT muscle from OMC stimulation were significantly faster than from LMC stimulation (**Figure 3B and D)**. The observed distribution of latencies from LMC stimulation to the CT muscle is consistent with a present, but sparse direct projection from the LMC region to Amb, that is dominated by indirect projections (Arriaga et al 2012).

**Figure 3.**
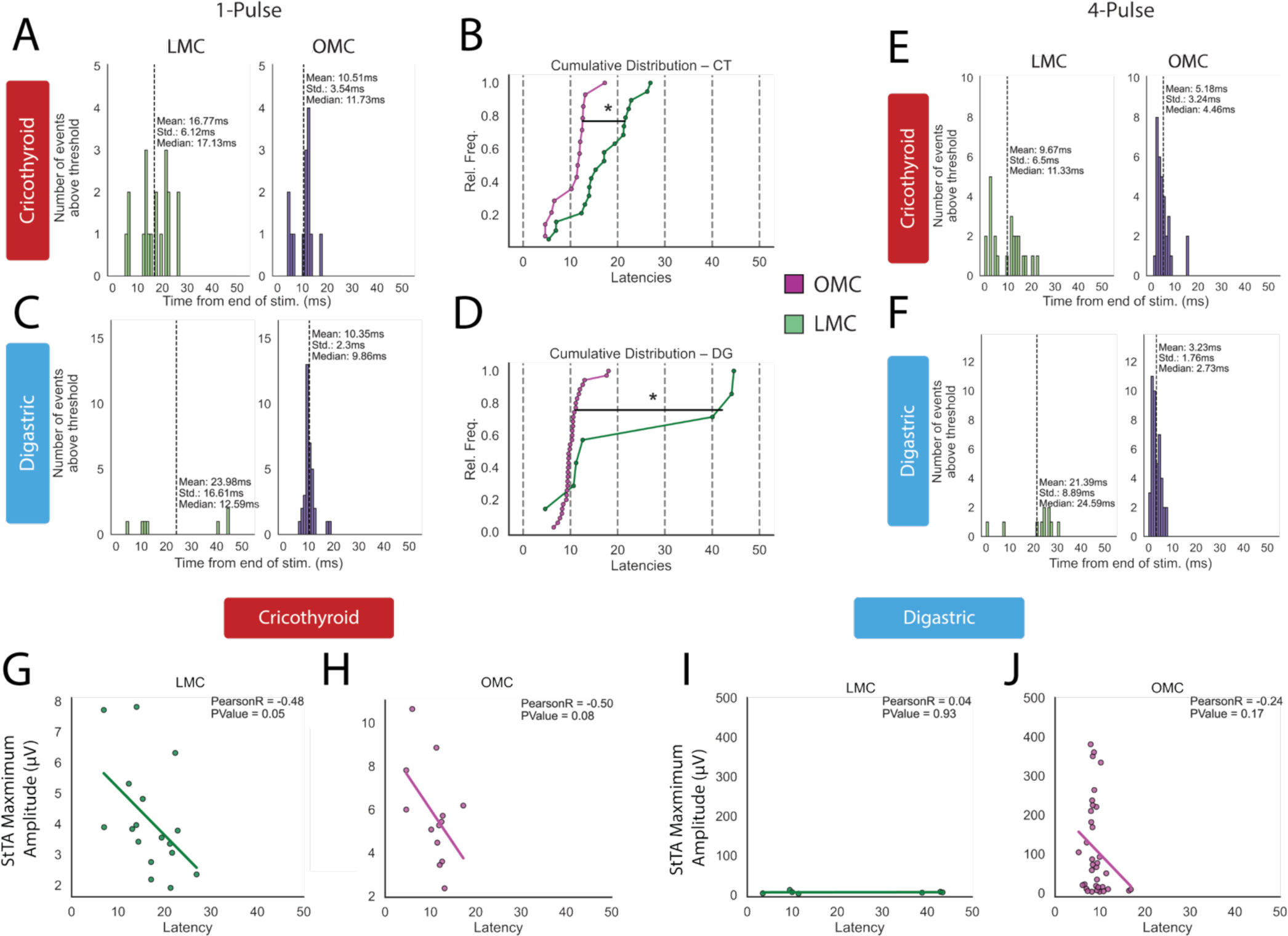
Latency and current-dependent properties of cortical to muscle responses. (A and B) Histogram (A) and cumulative distribution (B) of latencies calculated for the CT muscle StTAs for single-pulse cortical stimulations in LMC and OMC. OMC latencies are significantly earlier than in LMC (Kolmogorov-Smirnov test, p = 6.4×10^−4^). (C and D) Histogram (C) and cumulative distribution (D) of latencies calculated for the DG muscle for single-pulse stimulations in the LMC and OMC regions. OMC latencies to the muscles are significantly earlier than from LMC (Kolmogorov-Smirnov test, p = 0.031). (E and F) Histogram of latencies after four-pulse stimulation in LMC and OMC for the CT (E) and DG (F) muscles responses. (G and H) Relationship between latency and maximum amplitude for the CT muscle after stimulation in LMC (G) and OMC (H) with outlier values removed. (I and J) Relationship between latency and maximum amplitude for the DG muscle after stimulation in LMC (I) and OMC (J).

We also plotted the latency distributions from four-pulse stimulations, with the latency defined as the time of a response after the last pulse of the four-pulses. We observed generally similar properties as the single-pulse distributions, but with a higher proportion of cortical sites generating responses in each muscle. In the CT, there was a broad range of latencies from LMC stimulation (sites in n = 5 of 6 animals tested), while narrowly centered shorter latencies from OMC stimulation (sites in n = 6 of 7 animals tested) (**Figure 3E**). The 0 ms latencies are likely due to the first pulses leading to a response before the end of the full four-pulses. In the DG muscle, there were again fewer responses to LMC stimulation which tended to be slower (sites n = 3 of 6 animals tested), while all sites with OMC stimulations resulted in a muscle response, and the latencies were faster (sites in n = 7 of 7 animals tested; **Figure 3F**).

One might expect that with a mixed population of direct and indirect projections, the strength (e.g. amplitude) of the muscle response could be stronger in those that receive the direct cortical projection. To test this idea, we plotted the latency against the respective maximum amplitude (in µV) for each responsive StTA and determined any possible correlation. After removing three outliers with artifacts from LMC and one from OMC stimulation (**Figure S6A-C**), we found a negative correlation where the CT muscle’s fastest responses had the highest EMG contraction amplitude from LMC stimulation (**Figure 3G)**; a correlation with OMC stimulation and the CT muscle’s response approached significance, which is also tentative considering the responses were found in sites in just one animal (**Figure 3H**). In contrast, for the DG muscle, there were no significant correlations between muscle response latency and muscle response amplitude from either LMC or OMC stimulation (**Figure 3I and J**). DG muscle response amplitudes from LMC stimulation were very low around 10 µV (**Figure 3I**) and from OMC stimulation had a wide range from 10 to 400 µV (**Figure 3J**). These results are indicative of both direct and indirect projections, at least from LMC to CT.

We next tested whether latency or EMG muscle response amplitude could be affected by changes in the intensity of cortical stimulation, as in some sites muscle response amplitude increased with higher ICMS currents (**Figure S6D**). Due to the variability in the depth of anesthesia and current intensities used across mice, we performed this analysis per responsive site, rather than in aggregate. We selected sites that had two or more responsive StTAs in the muscles that were elicited from different current intensities. We then measured the slope of all correlations as a measure of the direction in which amplitude or latency could change with current intensity. We found that for the CT muscle response amplitude, there was no significant positive or negative correlation with different stimulation currents in either LMC or OMC (**Figure S6E**). For the DG muscle response amplitude, there was no significant correlation from stimulating LMC, but we found that stimulations in OMC had a significant number of slopes that were positive **(Figure S6F**), suggesting that increased current intensity in the OMC tended to increase the amplitude of the DG StTAs muscle response. The lack of change in amplitude for the CT muscle suggests we may already be inducing the respective regions maximal output to the muscle, even at lower stimulation intensities, whereas the DG muscle has a larger dynamic range. We performed a similar analysis with latency, and found that neither muscle showed significant increases or decreases with different stimulation currents in LMC or OMC (**Figure S6G and H**). The consistency in the response latency suggests we were not artificially inducing a short latency simply by having higher current intensities. Further, although direct projections exist, LMC activation of the CT muscle likely functions predominantly through the indirect pathway to Amb (as well as to trigeminal motor neurons for DG), whereas OMC activation of CT and DG muscles may be functioning through an undescribed direct connection to their respective motor neurons.

### Cortical neurons represent multiple muscles

In the standard homuncular model, subregions of M1 are seen as specialized for the activation of specific muscles, albeit with some inter-individual variability^36^. Thus, with our stimulations we were surprised by the amount of co-localized muscle activation from M1, particularly with the 4-pulse stimulations. Although the OMC region had been shown to control multiple muscles in mice^37^, we were particularly surprised by the overlap with the LMC region. To investigate the potential neural substrates for this co-activation, we injected the retrograde transsynaptic tracer pseudorabies-Bartha (PRV) labelled with different fluorophores (GFP and mCherry) into the CT and DG muscles of each mouse (**Figure 4A**), with the two fluorophores alternated between muscles across mice to control for possible differences in the virus transfection^40,41^. After a 96-hour incubation period^30,31^, we found a large overlap of transfected neurons in several subcortical regions such as the intermediate reticular formation and locus coeruleus (both single and co-labeled neurons; **Figure S7A**). Within M1, both CT- and DG-representing layer 5 neurons transfected with PRV were inter-mixed within the same cortical column (**Figure 4B**), in the same cortical space previously defined as LMC^30^. Neurons representing each muscle were adjacent to each other and some were co-labeled representing both muscles. We did not observe any transsynaptically labelled neurons in OMC. We counted the labelled neurons and found that 6.7-22.2% of the labelled layer 5 M1 neurons were co-labeled from CT and DG muscle injections (**Figure 4C and S7B**). This finding is consistent with a contemporaneous study, showing that in *S. teguina,* approximately 50% of CT- and DG-representing neurons were co-labeled^31^. The lower number of co-labeled CT and DG neurons in our study could be due to species differences. These findings are consistent with our ICMS experiments from the LMC region being able to activate both CT and DG muscles, although the latter less often.

**Figure 4.**
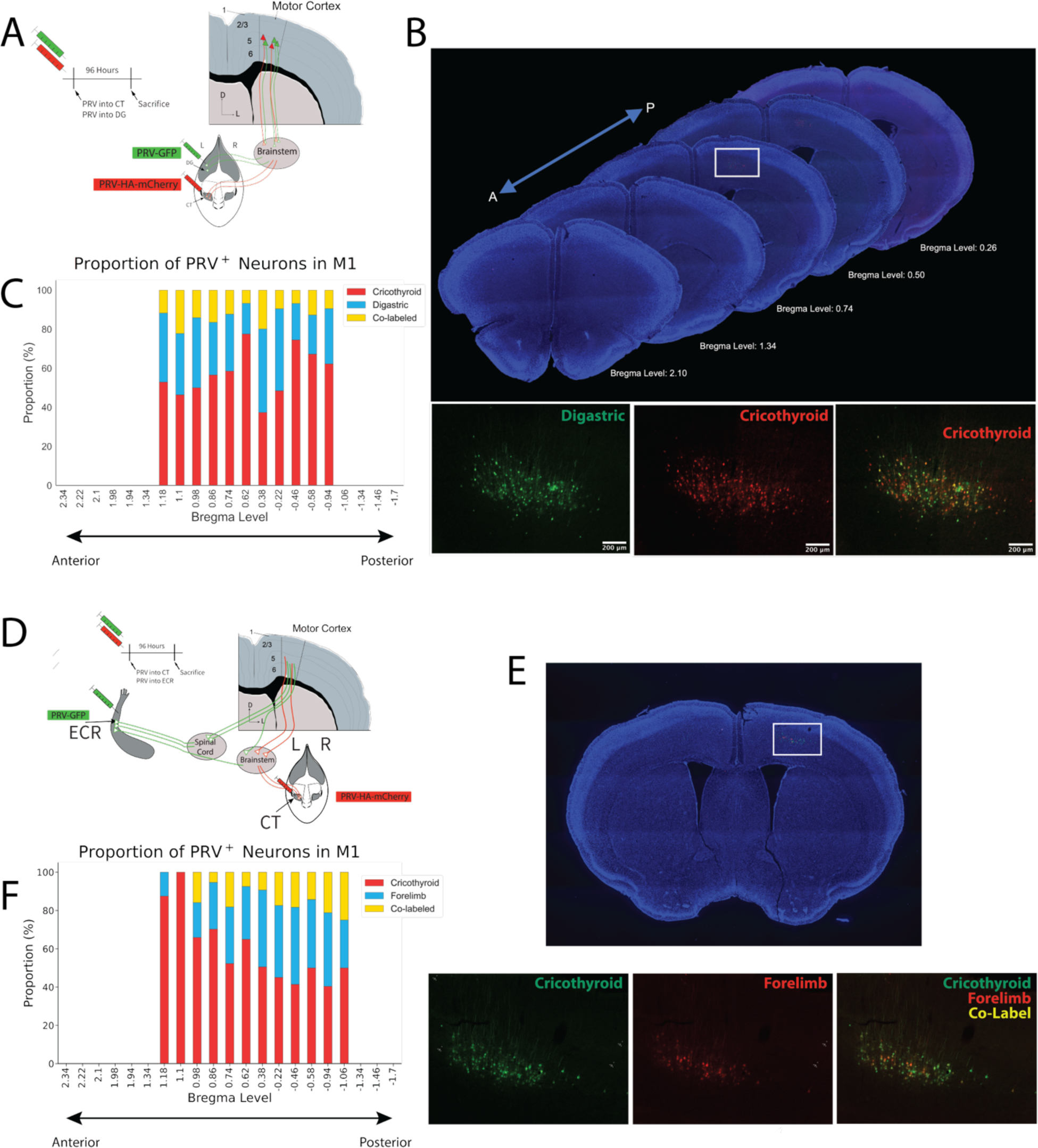
Retrograde Tracing with PRV from Two Pairs of Muscles. (A) Overview of experimental procedure to inject PRV encoding different fluorophores into CT and DG. Virus was alternated for each muscle across mice to balance potential differences in expression or tropism. (B) Coronal slices from a representative animal aligned anterior to posterior, and close-up of region outlined by the white box. Left, green neurons are transfected from injection into the DG; middle, red neurons are transfected from injection into the CT; right, merge depicting the single-labeled and double-labeled cells from the two muscles injected. Blue, DAPI stain (C) Mean proportion of cells counted to be either single- or double-labeled by PRV injected into CT and DG muscles. (A) Overview of experimental procedure to inject PRV encoding different fluorophores into CT and ECR (i.e. forelimb). Virus was alternated for each muscle across mice to balance potential differences in expression or tropism. (B) Coronal slice from a representative animal aligned anterior to posterior, and close-up of region outlined by the white box. Left, green neurons are backfilled from injection into the CT; middle, red neurons are backfilled from injection into the ECR; right, merge depicting the single-labeled and double-labeled cells from the two muscles injected. Blue, DAPI stain. (C) Mean proportion of cells counted to be either single- or double-labeled by PRV injected into CT and ECR muscles.

We also noted that the M1 region transsynaptically labeled from PRV injections in CT and DG muscles is similar to what has been described as the forelimb motor cortex by ICMS^36,42^ and transsynaptic tracing from the forelimbs^39,43^. Beyond the spatial coincidence, we wondered if this co-representation of muscles was unique to pairs of muscles that are synergistic for oral/facial movements, like CT and DG, or if this was a broader property of this part of mouse M1. To address this question, we again injected PRV with different fluorophores into the CT muscle and the ECR muscle of the forelimb (**Figure 4A**). We found that neurons representing CT and ECR muscles shared largely overlapping cortical space, as well as some individual neurons representing both muscles, like our findings in the CT and DG muscle injected mice (**Figure 4B**). The proportions of transsynaptic co-labeled cortical neurons from CT and ECR muscle injections ranged from 5.3-25% across M1 (**Figure 4C and S7C**). Again, we did not observe any transsynaptically labelled neurons in OMC.

We asked whether between the two co-labeling experiments, if there were differences in the of PRV-labeled layer 5 neuron distribution across the anterior-posterior axis of M1. The anterior-posterior distribution overlapped in both the relative precent (**Figure S7D**) and total number (**Figure S7E**) of transsynaptic single-labeled layer 5 CT-representing neurons in the CT-DG injections and CT-ECR injections; consistent with this, the cumulative distributions were not significantly different (Kolmogorov-Smirnov, p>0.05). However, there was a significant correlation in the proportion of single-labeled CT-representing neurons along the anterior-posterior axis of the CT-ECR injections (Pearson, p=0.0026). For the co-labeled layer 5 neurons, there was a decrease in the proportion of CT-DG-representing neurons and a significant increase of CT-ECR-representing neurons along the anterior-posterior axis (Pearson, p = 0.0261) while there was no significant difference in the cumulative distributions (**Figure S7F)**. These findings suggest that while there is some degree of “homuncular” organization in mouse M1 from anterior to posterior, much of this region of M1 has overlapping, gradient-like circuits around shared muscle representations, and perhaps behavior.

### OMC neurons form CM-like direct-projections to brainstem motor neurons

The lack of transsynaptically labelled PRV neurons in OMC from the CT, DG, or ECR muscles contradicted the finding of the short latency responses from stimulating some regions of OMC. Additionally, we are not aware of any study with PRV injected into muscles where OMC was successfully infected^44^. We sought an explanation and surmised that either: a) only an indirect projection exists from OMC to the muscle groups, but that pathway activation is much faster than the direct projection from LCM; or b) projections from OMC to the brainstem have different viral tropism for PRV than the LMC projections. To test for the presence of such projections that could be responsible for our short latency measurements with a different tracer, we injected the anterograde transsynaptic tracer AAV1-hSyn-Cre-WPRE^45,46^ into the OMC of Ai14 (CAG promoter tdTomato reporter) mice (**Figure 5A**). AAV1-transfected neurons expressing synapsin 1 will undergo CRE recombination to express tdTomato, and then transport down axons from OMC and jump one synapse to their brainstem targets^46^. We found sparse, but clearly present post-synaptic labelled neurons in the trigeminal motor nucleus (Mo5), which contains DG motor neurons (**Figure 5B**); we confirmed a previously described direct projections to the facial nucleus (7N; **Figure S8A**), which controls the whiskers^47,48^. We used a ChAT co-label to confirm that these post-synaptic neurons in Mo5 and 7N were in fact motor neurons (**Figure 5B and S8A**). We did not find any post-synaptically labeled neurons in the Amb (**Figure 5C**). In addition to these brainstem motor neuron targets, we found few OMC post-transsynaptic neurons in the lower layers of the LMC region (**Figure 5D**), but many in the neighboring primary somatosensory (S1) cortex. Beyond these regions, we found OMC targeted neurons in the ventrolateral striatum, thalamus (reticular thalamic nucleus, ventromedial nucleus, ventrolateral nucleus, and parafascicular nucleus) (**Figure S8B-D**; similar to Yang et al. ^49^), the substantia nigra, red nucleus, superior colliculus (SC) (**Figure S8E**), locus coeruleus, as well as numerous projections in the brainstem reticular formation, and oral part of the spinal trigeminal nucleus (SpVO). Since we did not find AAV1 label in Amb from OMC injections, we also injected AAV1 into the LMC of Ai14 mice to test if the previously identified direct projections could be identified with this method ^30^, but we did not find post-transsynaptically labeled neurons in Amb (**Figure S9**).

**Figure 5.**
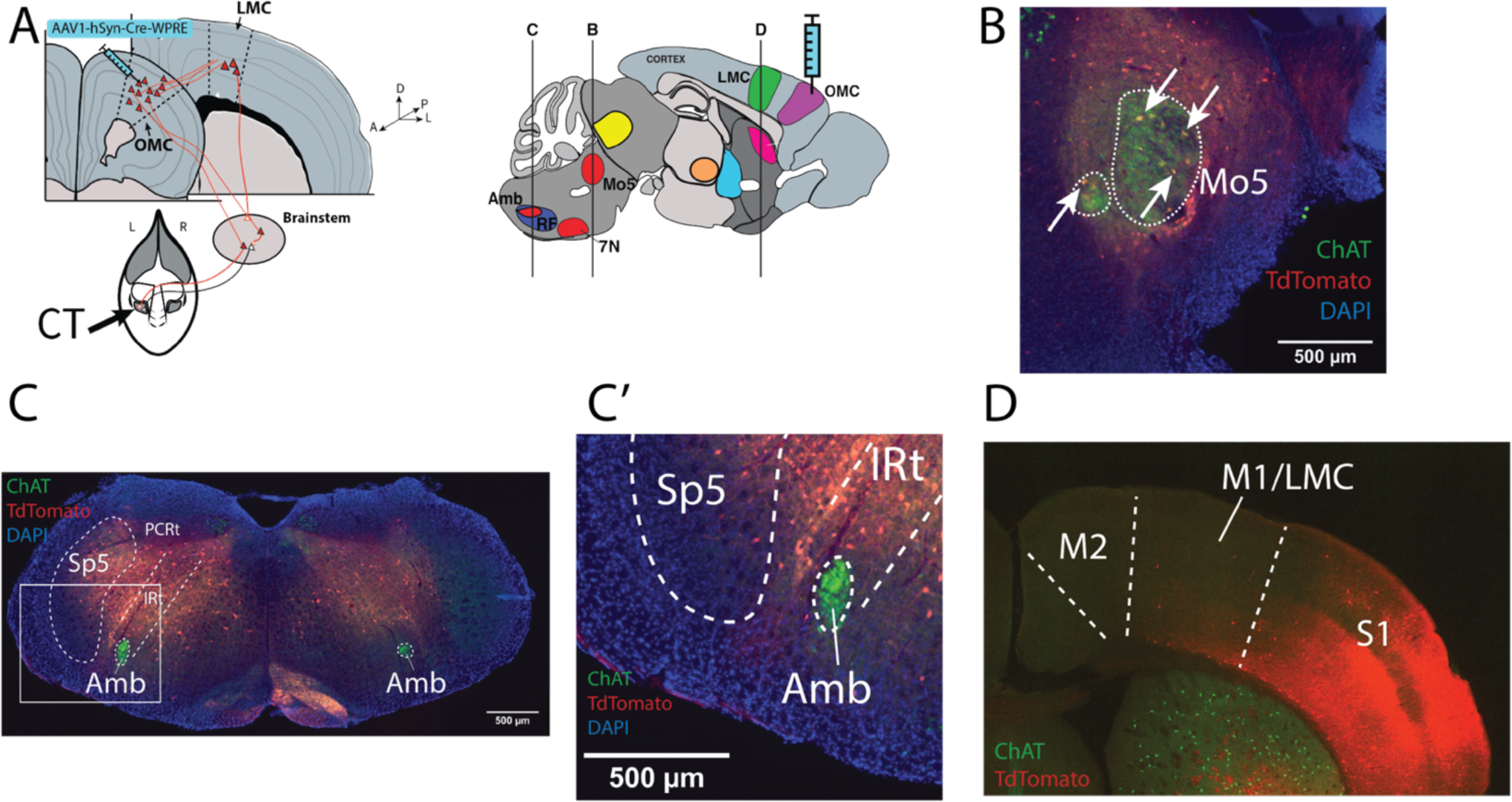
Efferent targets of OMC include direct projections to jaw motor neurons. (A) Left - overview of experimental procedure to inject AAV1 in Ai14 mice. Right – Sagittal diagram of mouse brain with lines representing approximate position of images in B, C, and D. (B) Example of post-transsynaptically labeled neurons (yellow and arrows) in the trigeminal motor nucleus (Mo5). Verified with ChAT double-label (green). (C) Example showing no post-transsynaptically labeled neurons seen in the nucleus ambiguous (Amb). A substantial number of neurons labeled in the surrounding reticular formation. Highlighted box is shown at higher magnification in C’. (D) Example showing some post-transsynaptically labeled neurons in the LMC region of M1 and S1. Many more projections seen in S1 lateral to LMC. Abbreviations: Mo5, motor trigeminal nucleus; Sp5, spinal trigeminal sensory nucleus; PCRt, parvicellular reticular formation; IRt, intermediate reticular formation; Amb, nucleus ambiguus; M2, secondary motor cortex; M1/LMC, primary motor cortex/laryngeal motor cortex; S1, primary somatosensory cortex.

To specifically visualize the axons from OMC to Mo5, we injected AAVretro-Cre-eGFP, a retrograde virus, into Mo5 and a AAV9-DiO-mCherry virus into the contralateral OMC (**Figure S10A**). Neurons expressing both viruses, through CRE recombination inside the same cell, will express mCherry, otherwise neurons projecting to Mo5 but not from OMC will only express eGFP. Axon terminals expressing eGFP from the OMC neurons were found within, and surrounding, the injected Mo5, as well as a few fibers in the contralateral Mo5 (**Figure S10B**). There were many eGFP-expressing neurons throughout the cortex, which may be due to virus spreading beyond Mo5 as the area around Mo5 receives dense motor and sensory input; but the layer 5 OMC neurons that project to Mo5 were clearly visible (**Figure S10C**). Interestingly, terminals from the mCherry expressing neurons were also found in many of the regions noted above from the anterograde transsynaptic-only injections. Some of these regions include the substantia nigra, red nucleus, and superior colliculus (**Figure S10D**). Other similarly targeted regions include the thalamus (ventromedial, ventrolateral, mediodorsal, and perifascicular nuclei), reticular formation, and SpVO. Among the cortico-cortico projections, we validated the strong projection from OMC to S1 and to ventrolateral striatum, but we did not identify any fibers to LMC (**Figure S10E-H**). Thus, the cell type that makes a weak projection from OMC to LMC in our AAV1 injections is a different population than the OMC population that projects to Mo5. These findings indicate that both LMC and OMC regions have sparse direct projections to different brainstem motor neuron groups (Amb and Mo5 respectively), but they have different tropism for different retrograde and anterograde transported viruses.

### OMC lesions alter vocal behavior

Lesions of LMC in humans, and the analogous RA in songbirds, leads to difficulties or loss in producing learned vocalizations^6,15,26^. Prior work from our laboratory demonstrated that lesioning LMC in mice leads to some degradation in the frequency modulation in USVs, but not loss of production of USVs^30^. In addition to their acoustic diversity, mouse USV songs exhibit a rich sequence diversity^50,51^. Since OMC stimulation was able to generate some vocal muscle contractions, we tested if OMC had a role in USVs, especially at the level of song structure and organization. We chemically lesioned OMC bilaterally in male mice using ibotenic acid (**Figure 6A**). Control mice were injected with saline. We elicited vocalizations from the male mice by introducing a female mouse into the recording chamber in 5 minute sessions. We performed baseline recordings of the male USVs in response to females 1 week prior to surgery then 2- and 3-weeks post-surgery. Vocal data were analyzed using Analysis of Mouse Vocal Communication (AMVOC)^52^.

**Figure 6.**
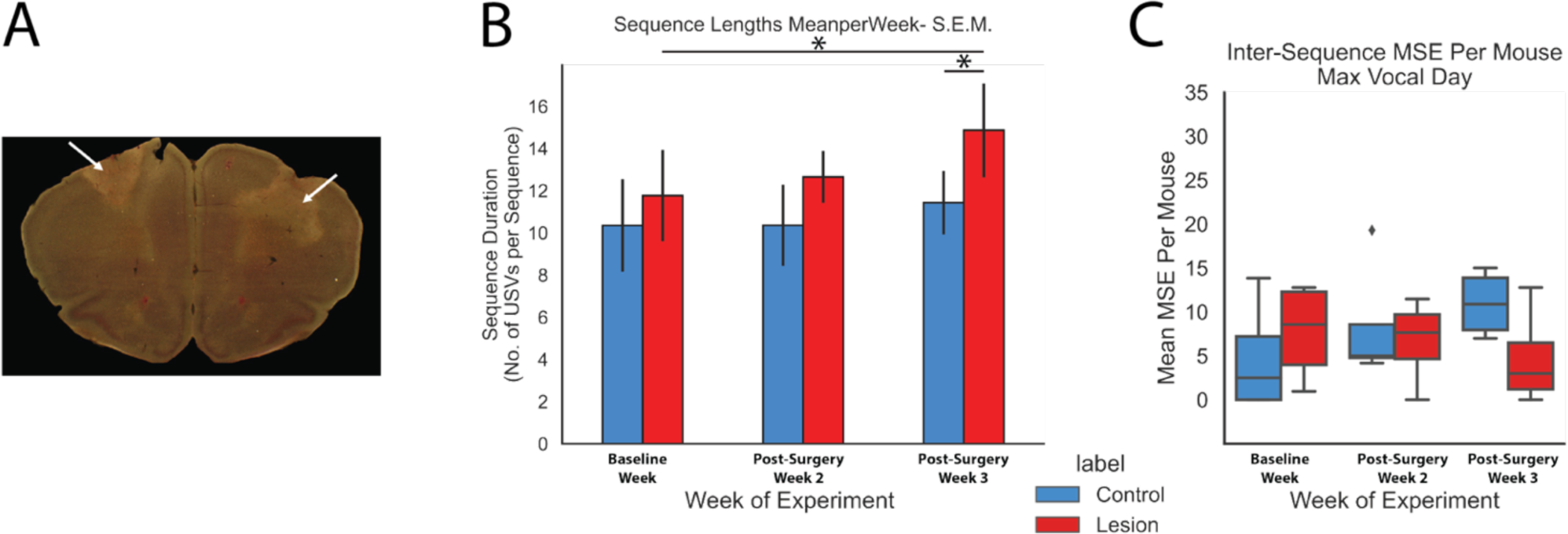
Impact of OMC lesion on USV production. (A) Representative darkfield Nissl image of a bilateral OMC lesion from ibotenic acid injections. (B) Length (in syllable number) of USV sequences before surgery, two weeks post-surgery, and three weeks post-surgery. There was a significant increase in the length of USV sequences in OMC lesioned mice (Paired t-test, p=0.012) and not in the control mice (Paired t-test, p>0.05). Lesioned mice had longer sequences compared to control mice at 3 weeks post-surgery (t-test, p=0.035). Bars indicate standard error of the mean. (C) Differences in syllable transitions between phrases measured by the mean squared error (MSE) between the transition matrices. Lower MSE indicates more similar matrices. There was no significant difference in inter-phrase transitions between weeks (Paired t-test, p>0.05).

We found that by 3-weeks post-surgery, lesioned mice had a greater number of syllables per vocal sequence phrase compared to baseline (Paired t-test, p = 0.012) and compared to controls (t-test, p=0.035; **Figure 6B**). Control mice exhibited no change in number of syllables per phrase across weeks (Paired t-test, p > 0.05). We considered a hypothesis that the mouse motor cortex may play a role in determining the changes in vocal syntax observed in different social contexts^53^. We performed an unsupervised classification of vocalizations with AMVOC^52^ into 5 syllable classes, and one additional class for silence (**Figure S11**). We calculated the mean squared error (MSE) between sequences (see Methods), but did not find any differences in MSE distance between weeks in either condition (**Figure 6C**; all paired t-test p > 0.05); there were also no differences between the groups at each time-point (Mann-Whitney p > 0.05). These results suggest that OMC modulates sequence length, but the specific syllable sequences are likely generated elsewhere in the vocal circuitry.

## DISCUSSION

This is the first report we are aware of that stimulation of M1 in a vocal non-learner generates vocal muscle contractions. Stimulation of the PRV-identified LMC region of M1 preferentially drives laryngeal muscle contractions, whereas the OMC region preferentially drives jaw muscle contractions. Given these preferences, both muscles may be activated from both regions of cortex. The response latency from LMC and OMC stimulation for both muscles and the viral tracing supports the presence of sparse direct cortical projections to the muscles’ motor neurons, such as those we identified in Mo5. There is also a large representational overlap in the cortical space, and some individual layer 5 neurons in the LMC region project to multiple muscle motor circuits. Unlike LMC’s impact on frequency modulation, lesioning OMC alters the number of syllables per phrase. Overall, these findings suggest a re-evaluation of prior theories about the role of M1 in vocal non-learning species and homuncular organization in rodents, which we discuss below.

### Cortical circuitry for vocalizations

Since the discovery that mouse USVs have complex song-like structures^51^, there has been great interest in understanding their mechanisms of vocal production and control. M1 is often understood to be critical for facilitating fine, volitional control over movements^4,18^, with the number of direct projections from M1 to motor neurons providing animals with a greater degree of fine control or dexterity over the targeted muscle groups^18^. After we first published our labs’ findings on the mouse laryngeal representation in M1^30^, there was still debate over the role of the cortex in controlling the mouse larynx. Several studies before and after demonstrated that many aspects of mice’s vocal repertoires and vocal features are genetically determined and heritable^54,55^. One study found that mutant mice lacking most of the cortex can still produce normal USVs—as well walk and eat on their own^56^, which led many to argue against any cortical involvement in this behavior. Later analyses of these mice’s vocalizations found subtle spectral changes, detectable with machine learning methods^57^. Our present data demonstrate that parts of mouse M1 can generate vocal muscle contractions, further supporting the hypothesis that mouse M1 plays some a role in vocal communication.

In humans, stimulation of vLMC and dLMC causes speech arrest and basic vocal production^11,58^. Instead, in non-human primates, stimulating a premotor region, area 6V, caused laryngeal muscle contraction, but no vocalizations^24,25^. Area 6V thus became the focus of vocal research in non-human primates, and was termed LMC in those studies^19^. Similar to our prior studies^30,59^, and this study, using retrograde transsynaptic viral tracers in mice, Cerkevich *et al.* ^33^ injected rabies virus into the laryngeal muscles of marmosets and macaques and demonstrated these primates also have a laryngeal representation in M1. Our study thus converges with lines of evidence from other species that cortical laryngeal representations are present in non-vocal learning species, including in rodents and non-human primates.

This laryngeal M1 representation in rodents may have gone unreported or overlooked as overt movement was used to define the function of areas^36,42,60^. In primates, laryngeal regions were described by direct palpation or a laryngoscope to detect laryngeal muscle movement upon cortical stimulation^24,61^, which can make it difficult to visually identify potentially small contractions. Instead, we used EMG recording of laryngeal muscles, which is more sensitive; we also had a transsynaptically-identified connected region of M1 that we could test as a candidate, independent of current stimulation and contractions of other muscles. We believe the combination of cortical stimulation in retrogradely identified regions provides a more unbiased approach to understanding the representational organization of M1. To our knowledge there have not been EMG measurements from the larynx in non-human primates during M1 stimulation. Considering these differences, it is important to know whether stimulating the transsynaptically-identified laryngeal connected M1 region in macaques and marmosets identified by Cerkevich *et al.* ^33^ leads to vocal muscle contractions measured with EMGs, and if so, what latency these contractions occur.

### ICMS estimates of synaptic order and monosynaptic connectivity

Prior to the advent of transsynaptic tracers, ICMS-EMG was used for determining the synaptic order of cortical control of motor neurons. In one classic study, when the forelimb region of macaque M1 was stimulated, the resultant latency of EMG in forelimb muscles was ∼10 ms^62^. Similarly, in recent studies in humans undergoing brain surgery, when the dorsal laryngeal motor cortex (dLMC) region was stimulated, laryngeal muscle EMG response latencies were in the range of 11-22 ms^12,63^. Other studies used transcranial magnetic stimulation (TMS) on the skull in the area of LMC (the exact area is difficult to define with TMS) which resulted in laryngeal contraction latencies of ∼10 ms^10,64^. Stimulations of area 6V in macaques result in a laryngeal contraction latency of 20 ms or longer^25^. In mice, when the number of direct projections to forelimb motor neurons was increased genetically by downregulating the PlxnA1 gene in the cortex, the mean latency of forelimb contractions from M1 stimulation was lowered from ∼13 ms in wild-type mice to ∼10 ms^65^. In our present study, stimulating the LMC region of mice, containing known direct-projecting layer 5 neurons, although latency of the laryngeal contraction has a mean of 24 ms, there is a range of sites with latencies from 10 ms to 45 ms. This range and distribution is supportive of a combination of sparse direct and predominantly indirect projections from the LMC region to Amb, and thus placing the mouse at a predicted intermediate position in the continuum hypothesis of vocal learning^34^.

We were surprised to find the OMC region is where we measured a higher proportion of faster (10 ms) latencies to the muscles than from LMC stimulation. Such a difference in latency between different regions of M1 for mouse forelimb responses has been reported from optogenetic stimulation of the cortex in^66^. In macaques, the hindlimb muscles are innervated by direct projecting layer 5 neurons, yet exhibit longer latency responses and show less facilitation from ICMS than their forelimb counterparts, hypothesized to be due to the lower synaptic strength^67^. One possibility to explain these differences from different parts of the M1 within species, is that the proportion of the direct cortical projections to one set of motor neurons (e.g. LMC to Amb) is less than the direct projections to another (e.g. OMC to Mo5). More experiments are needed to determine the impact from having different proportions of the faster monosynaptic and slower multisynaptic cortex-to-motor neuron connections.

### M1α and M1β somatotopy in M1

The motor homunculus organization theory postulates that localized parcels of cortex are dedicated to particular parts of the body, with a general body plan organized from foot to head along the medial to lateral axis in primate M1^68^. A similar medio-posterior to latero-anterior organization has been generally regarded to also exist in mice—with a dominance of the forelimb representation^36,42^. For this reason, we were surprised that both ICMS and transsynaptic tracing simultaneously from different muscles revealed largely overlapping larynx, jaw, and forelimb cortical areas^36,69,70^; this general area has also been shown to contain representations of the tongue^44^, vibrissae^71^. Due to the superimposed representations, we propose a distinct term for this area as M1 alpha (M1α), and we term the more anterolateral region of M1 that does not exhibit PRV labeling as M1 beta (M1β) (**Figure 7A**). Rather than referring to the cortical region by a muscle or body part reference, we propose to use LMC to denote the subpopulation of neurons representing the larynx and CFA to denote the subpopulation representing the forelimbs in M1α; OMC would denote the subpopulation of neurons representing the jaw, and other orofacial muscles, in M1β. This type of overlapping organization within M1α and within M1β could facilitate a wide variety of movements that use similar muscles but require distinct ensembles to perform an action. For example, forelimb behavior is known to differ along the anterolateral and posteriomedial cortical axis^72^.

**Figure 7.**
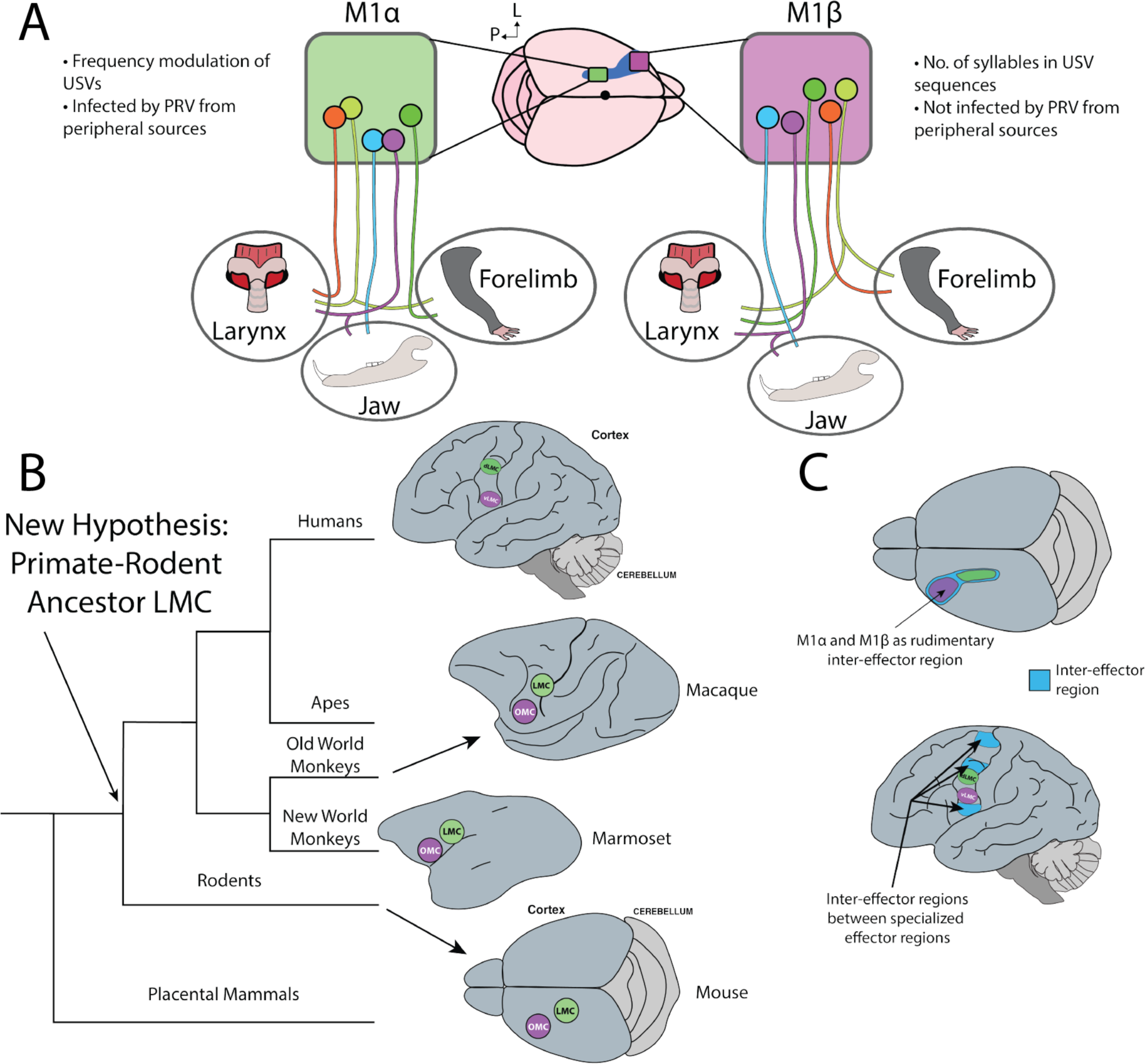
Hypotheses for the organization and evolution of vocal M1. (A) Schematic of proposed M1α and M1β organization of mouse M1. Individual neurons are represented as projecting to one or two different body parts that they may represent based on our data. We predict M1α and M1β are contributing differently to the overall output of M1 by representing different behaviors while representing similar muscles, similar to descriptions of action maps in primates ^82^. (B) Abbreviated phylogenetic tree of mammalian evolution with our new hypothesis that places the origin of an M1 LMC at the last common ancestor of primates and rodents, rather than an origin on the human lineage. Humans thus derived more distinctive and advanced LMC regions independently. (C) Potential comparison between the multi-representational organization of the proposed mouse M1α and M1β, human dLMC and vLMC, and their similarities to the inter-effector regions of the so-called “somato cognitive action network” in humans and primates described by Gordon et al. (2023).

Perhaps a hint of functional differences between the M1α and M1β regions is in the lesion findings. Lesions in M1α (i.e. LMC) result in changes in syllable frequency modulation^30^ and in M1β (i.e. OMC) result in changes in duration of sequences, similar to effects seen in *S. teguina* when the digastric representing region of M1β was cooled^32^. Further, a recent report conducted simultaneously with our study found that the duration of spiking activity in OMC scales with the duration of song in *S.* teguina ^73^. Other recent work has shown that OMC is also involved in oromanual motor behaviors, such as maintaining a food pellet near the mouth versus adjusting the pellet’s position^74^. Thus, M1β may be more involved with the broader duration of a behavior rather than fine-scale features within that behavior. The combined findings shown here and from others support a functional segregation of laryngeal and oral muscle representations between LMC and OMC that coordinates with each other during vocalizations.

We do not know if this co-representation of different muscles in the same part of cortex is specific to rodents, or more broadly applicable to mammals, including primates. If a species difference exist between primates and rodents, then the co-localization we proposed could be due to the lower density of neurons in rodent cortex compared to primates with similar brain mass^75^. With fewer neurons to dedicate to discrete representations, there could be more proximity between any two representations, or the necessity to multiplex, especially if they are synergistic muscles like opposing upper arm bicep and triceps muscles^65^. More anatomical data using multiple retrograde transsynaptic tracers injected into muscles of larger brained animals is necessary to better understand common principles of motor representations across mammalian species.

### Evolution of cortical control of vocal muscles

In contrast to the long-argued view that direct cortico-motor neuron projections are unique to primates, and that humans are further unique in having the LMC in M1 with a direct projection to Amb ^18,20^, based on the collective data across species^13,30,39^, we argue that a rudimentary M1-LMC region evolved at least as recently as the last common ancestor of rodents and primates (**Figure 7B**). Early in development, mice have many direct cortical projections to spinal cord motor neurons for the forelimb system, which are then pruned postnatally until they are effectively absent in adulthood^39,69,76^. With the genetic substrate for direct projections available in the ancestor of rodents and primates in M1α, we hypothesize that these transient, or in some cases remaining sparse, direct projections seen in rodents were “enhanced” or expanded upon in the primate lineage and in vocal learners for vocal motor neurons. We hypothesize that this then led to the more elaborate corticospinal system seen in human, with the dLMC and vLMC separating out of M1α and M1β, respectively, as distinct regions rather than a subpopulation. As with the somatotopy argument above, the so-called “corticalization” of movement (i.e. greater dependance on direct projections from cortex) may also be due to changes in neuron count and density^75,77^. The degree to which a species has these direct projections in their vocal circuitry would partially dictate their position along a vocal learning continuum^3,29,30^.

Under the previous assumption that non-human primates did not have an LMC in M1, several hypotheses were developed to explain how human LMC may have evolved. One hypothesis posited that the premotor cortex area 6V is the ancestral vocal region in primates and that in humans it migrated into M1 to become what is now known as vLMC ^27^. This was followed by a duplication of vLMC that migrated dorsally in M1 to become dLMC in humans^5^. An alternative hypothesis based on our findings here in mice, is that there were two rudimentary laryngeal representations phylogenetically older than primates (e.g. LMC in M1α and OMC in M1β). The relative position of mouse LMC, human dLMC, and monkey M1 LMC regions^33^ indicates that they could be homologous rather than the human dLMC arising completely *de novo*. Additionally, like human dLMC^12^, mouse LMC may function in pitch modulation of vocalizations^12,30^. Primate vLMC and mouse OMC have similar representational overlaps of the orofacial and laryngeal muscles^25,37,61^. Non-human primate Area 6V, in this view, we propose would be the premotor LMC region anterior to vLMC in M1. In this hypothesis, the biggest difference between mice and humans would be a segregation of the human LMC regions from other M1 functions, and the strongly enhanced direct projection to vocal motor neurons.

This hypothesis begs the question: is this dual cortical representation specific to vocalization across species, or, as we see in mice, does it contain other functions. In a recent study by Gordon et al^78^, primate M1 (including humans) was proposed to be composed of dual mirror image representations of three body segments (face area, upper limbs and, and lower limbs), termed effector M1 regions, and each set of effector regions separated by inter-effector regions that represent movements of many body parts. Taking this view into consideration and our findings in mice, we propose that mouse M1α and M1#x03B2; may represent ancestral dual effector regions observed by Gordon et al., which in humans contains the more advanced dLMC and vLMC (**Figure 7C**). An alternative is that mouse M1, in its entirety, is instead like an inter-effector region with overlapping gradients of representation, which remained rudimentary and undifferentiated in rodents while the primate linage derived the more specific representations of effector regions through the presence of increased neuron density and direct projections. To test any of these hypotheses, developmental, molecular, and genetic data must be collected and compared between these multiple laryngeal and other cortical representations across species.

## Acknowledgments

We thank many members of the Jarvis lab for their comments throughout the progress of these experiments, in particular: J. Lomax Boyd, Matthew Davenport, and Peter F. Schade. We thank John Kalambogias and Rodrigo Alonso for advice on the ICMS and EMG procedures, and Lisa Pomeranz for assisting us with the PRV used in this study. We thank Priya Rajasethupathy and A. James Hudspeth for their advice on these experiments, and Yutaka Yoshida for comments and discussion. This work was supported by a Keck Foundation award, an NIH Transformative Research Award R01 OD028000, and HHMI funds to E.D.J.; and an HHMI Gilliam Fellowship to C.D.M.V.

## Contributions

C.D.M.V and E.D.J. designed and conceived of experiments. C.D.M.V, E.N.W. performed ICMS and EMG experiments. C.D.M.V. analyzed EMG data. C.D.M.V. and R.K.A. performed surgeries and analyzed PRV experiments. C.D.M.V., R.K.A., and H.B. performed histology. C.D.M.V. performed lesions and collected vocalization data. C.D.M.V. and C.B. analyzed vocalization data. T.G. and E.D.J. supervised vocal analysis. C.D.M.V. and E.D.J. wrote the manuscript.

## Methods

### Animals

Adult (greater than 8-weeks old) male and female C57B6/J and Ai14 mice were used for this study (Strain #s 000664 and 007914, respectively, Jackson Laboratories, Bar Harbor, ME). Males and one female were used for stimulation experiments; both males and females were used for tracing experiments in approximately equal numbers; only males were used for lesion experiments; females were used as stimuli to induce male USV song production. Mice were housed socially with the same sex (except for eliciting USVs) on a 12-hour light cycle with *ad libitum* food and water. All animal protocols were approved by The Rockefeller University IACUC committee.

## ICMS and EMG

### Surgery

Prior to the surgical and experimental procedures, mice were anesthetized using a ketamine/xylazine cocktail (100mg/kg and 20mg/kg, respectively) given at 0.1ml/10g. Mice were anesthetized until they were not responsive to a toe pinch. Throughout the procedure, mice were supplemented with ketamine as needed. The mouse was placed supine on the surgical space. To access the DG muscle of the jaw, a small incision was made at the midline of the neck to expose the trachea and the anterior digastric muscle. Subsequently, to access the CT muscle of the larynx, the sternohyoid muscle was cut at the midline and held aside with retractors. For forelimb EMGs, the mouse was place prone on the surgical area and an incision was made along the length of the forearm to access the extensor carpi radialis (ECR) muscle. A pair of wires, forming a bipolar EMG electrode, were inserted into the CT and secured with a small dab of Vetbond^TM^ Tissue Adhesive (3M). Next, a second pair of electrodes was inserted into the belly of the DG. We used a 0.0026” diameter wire for DG and ECR, and a 0.0015” diameter wire for the CT as the larger diameter could not be easily accommodated in the CT. The tips of the electrodes were deinsulated. All EMG electrodes were made from formvar insulated nichrome wire (A-M Systems, Sequim, WA). Electrode wires were threaded through a pulled glass pipette (Sutter Instruments, Model P-1000). A slight bend was made on the inserted end of the electrode to provide a fishhook that would then remain attached to the muscle tissue.

After EMG wires were inserted into their respective muscles, the incision was closed using Vetbond^TM^. The mouse was the rotated to a prone position and placed in the stereotaxic frame. The fur on the skull was removed and an incision was made along the midline of the skull. Using a dental drill, a large (∼3mmx3mm) craniotomy window was made to expose the surface of the cortex contralateral to the muscles containing the electrodes. The dura was removed, and the surface of the brain was maintained wet using saline throughout the procedure.

### Stimulations and EMG Recordings

We performed cortical penetrations at intervals of ∼250 µm, with deviations to this dimension to avoid blood vessels. Penetrations were made non-sequentially so as to avoid biasing measurements due to the effects of anesthesia. There was an overall emphasis on the LMC and OMC regions in our sampling. Coordinates labeled as LMC were between −0.5-1.0 mm anteroposterior (AP) and 1-1.5 mm mediolateral (ML). These were based on PRV tracing ^30^. Coordinates labeled as OMC were between 1.5mm lateral and the lateral edge of the craniotomy, and approximately 1.8 anterior to the anterior edge of the craniotomy. These were based on responsive sites obtained for DG by^79^ and descriptions of the ALM regions ^44^. At each stimulation site, microelectrodes (50 µm diameter; FHC, Bowdoin, ME) were lowered to the approximate depth of layer 5B (∼800 µm for LMC and ∼900 µm for OMC—due to the increased curvature of the cortex). For stimulations we used an isolated pulse stimulator (Model 2100, A-M Systems), using current injections between 30-525 µA. For single-pulse ICMS we used one 0.2 ms biphasic pulse provided at a 2 Hz rate. For four-pulse ICMS we used an 8.5 ms duration, 0.2 ms biphasic pulse, 2.3 ms inter-pulse interval, provided at 1 Hz. Each round of stimulations lasted 30 seconds and constituted one recording and the same current was maintained throughout each 30s period. Our parameters resulted in 60 single-pulse stimulations per round and 30 four-pulse stimulations per round. All seven mice used received at least one test of both single-pulse and four-pulse stimulation.

EMGs were recorded using a differential amplifier (Model 1800, A-M Systems). All recordings had a system notch filter and signals were bandpass filtered (10 Hz – 5 kHz). Amplifier output was split between an oscilloscope, a speaker, and a recording DAQ (USB-231, Measurement Computing). For six mice, data from the amplifier was captured at 15kHz with the DAQ and streamed to the DAQami^TM^ (Measurement Computing) software on a nearby laptop. One mouse had signals recorded at 20 kHz per channel. These recordings were then downsampled to 15 kHz to be comparable with the other recordings.

### Data Processing

Signals from each recording were filtered with a 4^th^-order 1kHz high-pass Butterworth filter in the forward direction using the sosfilt function in sci-py Python library. After filtering, signals were full-wave rectified by taking the absolute value of the measured voltages. To get a reliable quantification for stimulus-responses from each cortical site and muscle combination and reduce breathing signal contamination in the CT muscle, for each current tested we calculated the mean of each round (30 seconds) of 60 (single-pulse) or 30 (four-pulse) simulations at a given cortical site within an animal, termed the stimulus triggered average (StTA). One StTA was computed for each level of stimulation at a given site in an individual mouse. For example, if a current was tested more than once at a site, there would only be one StTA.

All StTAs were manually inspected to eliminate obvious artifacts, such as increased, non-replicable, muscle EMG activity after 50 ms. We only considered StTAs that had putative responses to at least two different stimulation currents that were not 500 µA or above. To determine response latencies, for each putative positive StTA, we first determined the mean and standard deviation (SD) of the 30 ms preceding the stimulus artifact. The threshold was then set at 2 SD above the post-stimulation mean. In the post-stimulus period, if the measured voltage in the post-stimulation period crossed the threshold for more than ∼0.5 ms, then we considered that to be an EMG contraction. The first point of the EMG response to cross the threshold was used to measure the latency of the response for that cortical site-muscle combination. The latency for responses was determined based on the time between the end of the stimulus and the start of the response as calculated above. A buffer of a few ms was added in the post-stimulus period so that artifacts that lagged before the stimulus would not be counted as positive responses. Maximum amplitude was calculated as the highest voltage value measured in a muscle after the onset of the EMG.

All data were processed using Python and other freely available libraries developed for Python, including: numpy, pandas, sci-py, matplotlib, and seaborn. Correlations were determined using the scipy-py library’s Pearson function. Libraries can be downloaded with their respective commands from the Python Package Index (PyPI).

### UMAP and K-Means Clustering

All StTAs from mice were used to produce low-dimensional representations using Uniform Manifold Approximation (UMAP). First, we normalized each StTA with a maximum absolute scaler using the sci-kit learn library. These outputs were then fed to the UMAP with the following parameters (minimum distance=0.3, n-neighbors=50, distance metric=Canberra, random state=20). UMAP was performed in Python using the umap-learn library. For K-Means clustering, we performed the clustering on the normalized StTAs, also using the sci-kit learn library. We performed the K-Means clustering using 4 clusters as that was the number of combinations we had for each muscle (CT and DG) and each region (LMC and OMC) combination. Points on the UMAP projection were then colored by their respective K-Means cluster label.

## Viral injections

All viral injections were performed using the Nanoject III (Drummond Scientific). Injection needles were made from borosilicate glass and pulled using a Model P-1000 puller. The needles were backfilled with mineral oil.

### Pseudorabies-Bartha Virus (PRV) injections

We used two transsynaptic PRV constructs, PRV-HA-mCherry/PRV-lp298 (titre: 6.8×10^8 pfu/ml) ^41^ and PRV-GFP/PRV-lp297 (titre: 6.6×10^8 pfu/ml) ^40^, provided by Jeffrey Friedman (Laboratory of Molecular Genetics, The Rockefeller University). For these experiments, both constructs were injected into each mouse, one construct per muscle. We alternated which construct was injected for each muscle between mice.

Injection procedures were similar to those performed previously by our lab ^30,80^. Mice were anesthetized with 1-2% isoflurane with oxygen. Mice were then placed supine with a nose cone continuously providing anesthesia to the mouse. The fur at the site of the incision was removed and betadine was applied to the skin. A midline incision was made at the level of the neck, from between the shoulders up to the interramal whiskers to expose the larynx and digastric muscles in one incision. The skin and fat were separated and maintained in position using retractors. For the DG muscle, injections were made without any further procedures. To access the CT muscle, as with the EMG surgery, the sternohyoid muscle was cut at the midline and held apart with retractors. Injections are then made on the exposed muscle. As the CT is small and mice exhibit deep breathing in the supine position under anesthesia, care was taken to prevent the injection needle from penetrating below the muscle.

For the CT muscle, we injected 50 nl of PRV four times at four different sites at a rate of 15 nl/s. For the DG muscle, we injected 125 nl five times at four sites at 15 nl/s. The difference in volume was selected to account for the difference in the size of the muscles. After all the injections were completed, the separated tissues were closed and secured with Vetbond™. Mice were then administered bupivicane subcutaneously (0.25 mg/ml) and meloxicam (2-5 mg/kg). Post-operatively, mice were placed in a recovery cage placed on a heating pad and observed until they ambulated without issue. Once recovered, the mice were returned to their home cages. We used 5 mice for CT and DG co-injections of PRV.

For ECR injections, we performed them in combination with the CT. We used similar procedures as above. After the neck incision was made for the CT injection, the incision was closed as above. Mice were then laid prone in the surgical area. An incision was made on the skin of the forelimb on the dorsal side of the limb. We injected 125 nl five times at four sites at 15 nl/s, as in the DG muscle. The incision on the arm was secured with Vetbond™ and post-operative care was performed as above. We used 3 mice for CT and ECR co-injections of PRV.

### Adeno-associated viruses (AAVs)

Mice were anesthetized using 1-2% isoflurane with oxygen. After being placed in the stereotax, the fur on the scalp was removed. Betadine was applied to the exposed skin. A midline incision was made to expose the skull. The necessary number of craniotomies were made using a dental drill. For cortical injections, the needle was lowered into the brain up to 100 µm past the target and retracted into the target depth. For brainstem injections, the needle was lowered directly to the target depth. After injections were completed, the needle was left in the brain for 5 minutes to prevent injected medium from backflowing out of the brain onto the surface. The needle was retracted, and the skull scalp was closed at the midline with Vetbond™. OMC injections were made at +2 mm ML, +2.2 mm AP, and 0.9 mm below the surface. LMC injections were made at +1.4 mm ML, 0.3 mm AP, and 0.8 mm below the surface. For Mo5 injections, we injected at +1.4 mm ML, −6 mm AP, at an angle of 17° and a travel distance of 4.2 mm from the surface. For anterograde transsynaptic tracing we used AAV1-hSyn-Cre-WPRE. For combined OMC and Mo5 injections we used AAV5-DIO-mCherry and AAVrg-GFP-Cre, respectively. We used 6 mice for AAV1-hSyn-Cre-WPRE injections into OMC, and 4 mice for AAV1-hSyn-Cre-WPRE injections into LMC. We used 3 mice for dual injections of AAVrg-GFP-Cre in Mo5 and AAV5-DIO-mCherry in OMC.

## Perfusion and cryosectioning

Animals were transcardially perfused using cold 1X PBS and cold 4% paraformaldehyde (PFA) in 1X PBS. Extracted brains were post-fixed in 4% PFA overnight at 4°C. Brains were cryoprotected in 10% sucrose in 1X PBS solution overnight at 4° C, followed by 30% sucrose in 1X PBS at 4° C until the brains sunk. Brains were then placed in tissue molds and frozen with Tissue Plus^®^ O.C.T. Compound (Fisher Scientific). After freezing, the brains were cryosectioned (Leica CM 1950) at 50 µm and stored in 1X PBS at 4° C until used. Some sectioned brains were stored in 1X PBS with 0.01% sodium azide.

## Histology

For histology, we selected every 6^th^ section from the brain, unless otherwise noted. At this interval sections were separated by ∼300 µm from one another.

### Immunohistochemistry

All immunohistochemistry was performed with free-floating sections. Sections were washed in 1% Tween-20 in PBS for 3×15 min. Sections were then blocked in 1% bovine serum albumin (BSA) with 0.3% Triton-X in PBS for 2 hours. Following blocking, the sections were incubated in primary antibody with 1% BSA, 0.1% Triton-X in PBS overnight. On the following day, the sections were washed with 1% Tween-20 in PBS for 3×15 min and placed in the secondary antibody with 1% BSA and 1% Tween-20 in PBS for ∼2-4 hours protected from light. Sections were then washed with 1% Tween-20 in PBS for 3×15 min. Sections were mounted using VECTASHIELD^®^ Antifade Mounting Medium with DAPI (Vector Laboratories) and imaged on a fluorescent microscope (Olympus BX61 or Olympus MV×10). Images post-processed in ImageJ. We performed antibody labeling on the following targets: green fluorescent protein (GFP), tdTomato and red fluorescent protein (RFP; same antibody used), and choline acetyl transferase (ChAT). The primary and secondary antibodies used and their respective dilutions are listed in Table 1.

**Table 1:** 1° & 2° Antibodies used for immunolabeling 50 um adult mouse brain sections.

| 1° Ab | 1° Ab Concentration |
| --- | --- |
| Rockland Chicken anti GFP | 1:1000 |
| Rockland Rabbit anti RFP | 1:1000 |
| Rockland Goat anti ChAT | 1:100 |
| Rockland Chicken anti GFP | 1:1000 |
| Rockland Goat anti GFP | 1:1000 |

| 2° Ab | 2° Ab Concentration |
| --- | --- |
| Invitrogen 488 Goat anti Chicken | 1:500 |
| Invitrogen 568 Donkey anti Rabbit | 1:500 |
| Invitrogen 594 Goat anti Chicken | 1:500 |
| Invitrogen 488 Donkey anti Goat | 1:500 |
| Invitrogen 594 Donkey anti Rabbit | 1:500 |

### Nissl

OMC lesions were confirmed on naïve tissue or using Nissl staining. Briefly, sections were washed in 0.1M PB for 10 min, followed by 2 min in ddH_2_O. Sections were then placed in cresyl violet staining solution for 15-20 min. After staining, sections were placed in ddH_2_O for 1 min, followed by an ethanol gradient of 50% EtOH and 70% EtOH for 8 min each. Next sections were placed in 95% EtOH for ∼1 min with a few drops of acetic acid. Sections were moved to 100% EtOH for 2 min, and a separate fresh 100% EtOH for another 8 min. Sections were then washed in Histo-Clear^®^ (National Diagnostics) for 8 min 3 times and coverslipped using DPX Mounting Medium (Sigma-Aldrich).

### Stereology

To perform cell counts of the co-PRV injected brains (see above), we used the “Cell Counter” plugin in ImageJ. Green, red, and yellow (co-labeled) cell bodies were counted by adding these parameters as ‘counters’ in the FIJI cell counter plugin. Cell count outputs were analyzed and visualized using custom scripts written in python.

## Cortical lesions

### Vocal behavior

At 8 weeks of age, group-housed (up to 4) male mice were sexually socialized by placing an adult female in their home cage overnight at three days before the start of the experiment. Female-directed recordings were performed as previously described, using similar equipment ^53^. This equipment includes a cooler for sound isolation (Igloo^®^). We recorded vocalizations using UltraSoundGateCM16/CMPA ultrasonic microphones which were connected to an Ultrasound Gate amplifier, and digitally recorded using AvisoftRecorderUSG software (Sampling frequency: 250 kHz; FFT-length: 1024 points; 16-bits). Microphones, amplifier, and software are all from Avisoft Bioacoustics^®^ (Berlin, Germany). A set of pre-surgical baseline recordings were made for each mouse over three consecutive days. On the fourth day, surgeries were performed on the mice. Mice were singly housed for 4-5 days to allow for recovery from the surgery. After this recovery period, mice were placed back with their original cage mates for the remainder of the experiment. Three consecutive days of recording were repeated at 2 weeks and at 3 weeks post-surgery. Throughout the experiment, care was taken not to expose a male mouse to any one female more than once. At the end of the last recording session the mice were euthanized, transcardially perfused, and brains were extracted for histological processing.

### Surgery

Surgery and injections were performed with similar procedures as described above. Mice were injected bilaterally in OMC (+2 mm Lateral, and +2.2 mm Anterior) with a total of either 1% ibotenic acid or with saline as a control. Injections were performed at 1200µm below the cortical surface and 800 µm with 120 nl being injected at each level. We injected 40 nl 3 times, at 1 nl/s, with 60 seconds between each injection. After the second set of injections, the needle was removed from the brain. Once all injections were completed, the scalp was closed, and post-operative care was provided.

### Acoustic analysis

Audio files were processed using <u>Analysis of Mouse Vocal Communication</u> (AMVOC) software we developed ^52^. Audio files were processed using AMVOC and clustered into 5 different categories, and a 6th category of silence syllable (defined as a period of length greater than 250 ms with no vocalizations). Silent “syllables” were used for determining syllable classes that begin and end phrases. We used these silent periods to separate each set of consecutive vocalizations into sequences (Chabout et al., 2015). A sequence was defined by being composed of 2 or more syllables.

### Syntax analysis

For the syntax analysis, we did not include sessions where mice vocalized <20 USVs in 5 min. As found in (Chabout et al., 2015), this type of limited data skewed our syntax results, and so we performed our syntax analyses on the maximum vocalizing day for each week per mouse. To model the syllable transitions within sequences, we performed our analysis similar to what we described previously ^81^. Briefly, we first used AMVOC’s unsupervised classification (5 syllable types and 6^th^ silent “syllable”, **Figure S11**). We then represented transitions between each syllable type in a two-dimensional matrix where the rows and columns represent the different syllable categories, and each element equals the absolute probability of the corresponding transition. To examine the similarity between the transitions of two song sequences we applied an elementwise Mean Square Error (MSE) criterion to the corresponding transition matrices with values close to zero implying more similar transitions of syllables. The MSE between each sequence is calculated, and an overall mean MSE is calculated for each mouse in each week. We used this method to examine the changes in temporal structure between the first week and the next two weeks after the surgery. A paired t-test was applied to the same animals across weeks, and a Mann-Whitney U test between the control and lesion groups.

---

**Supplementary Figure 1.**
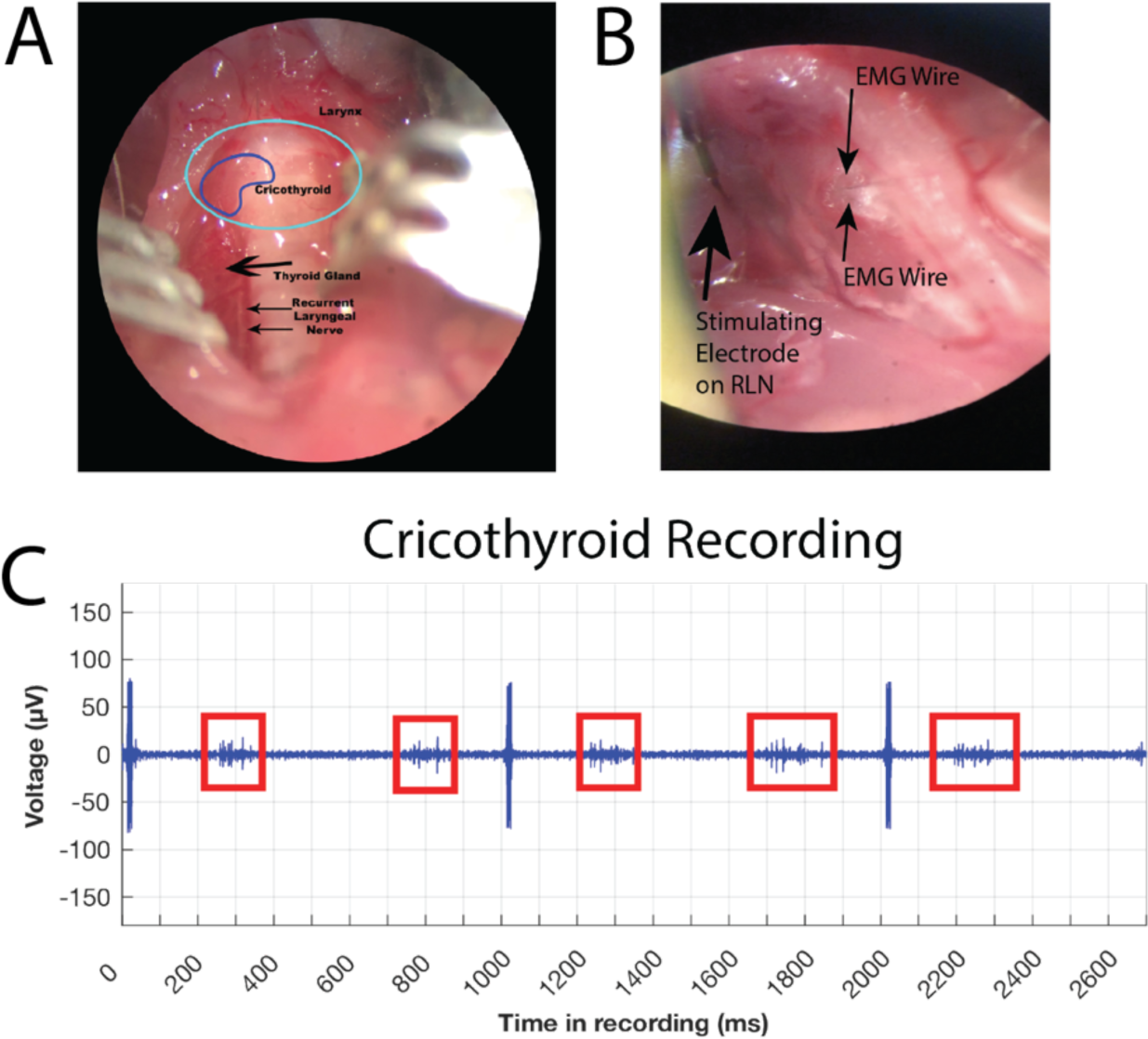
Acquiring EMG signal from the cricothyroid muscle. (A) Photograph depicting the anatomy of the neck where the larynx is located (light blue circle). The cricothyroid (CT) is outlined in dark blue. The sternohyoid muscle has been retracted to expose the trachea and underlying structures. (B) Photograph depicting a pair of fine wires inserted into the CT muscle. The wires have been affixed to the muscle by a light application of dermal glue (see Methods). (C) Unfiltered and unrectified EMG recording from the CT muscle during ICMS. Contractions from the CT muscle during breathing are highlighted (red). Arrows indicate stimulus artifact from ICMS.

**Supplementary Figure 2.**
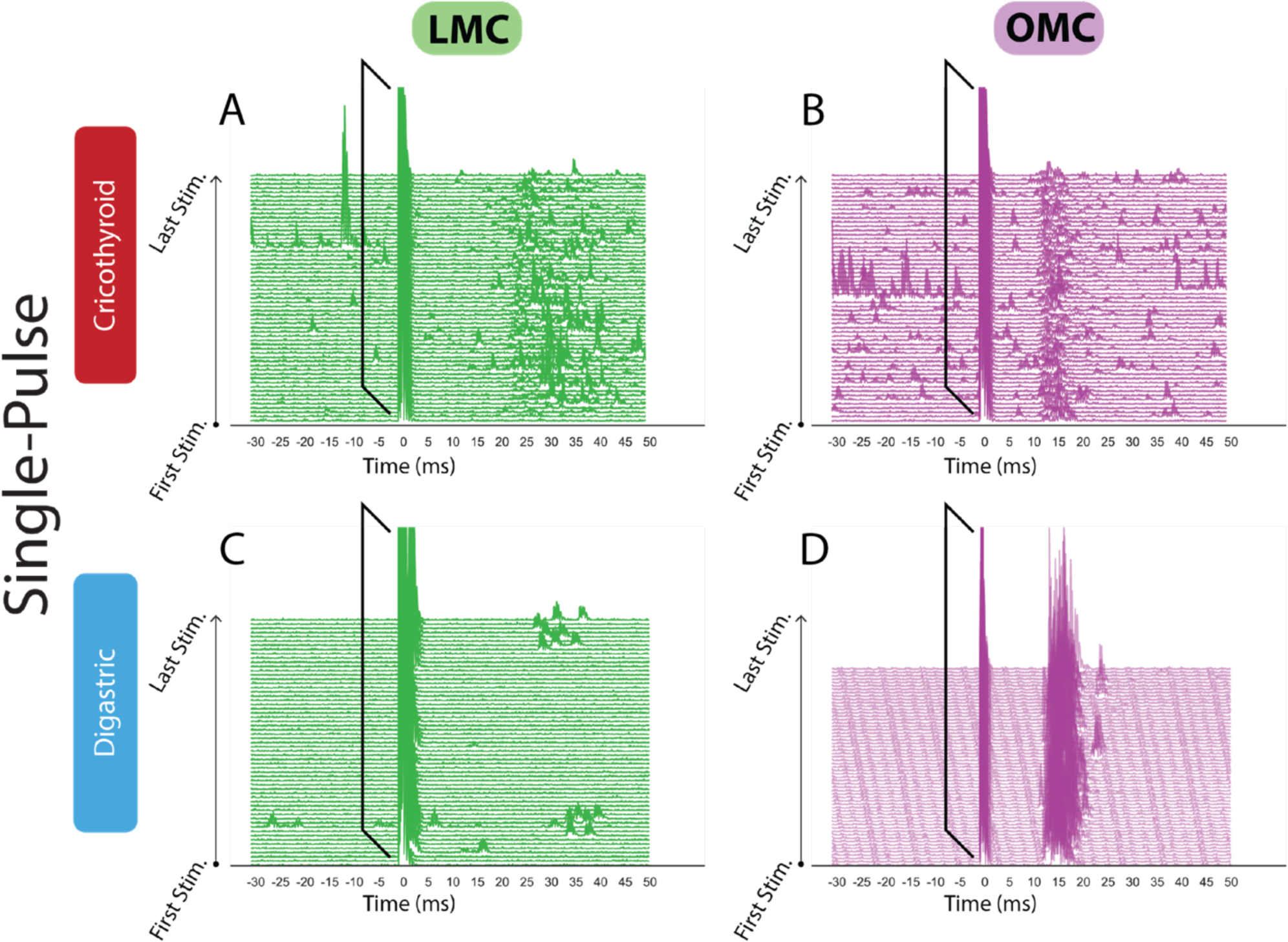
Example traces after single-pulse stimulation – Data from. Figure 1 (A-D) Individual traces from a single round of stimulations shown in Figure 1A-D, respectively. Brackets indicate stimulus artifacts.

**Supplementary Figure 3.**
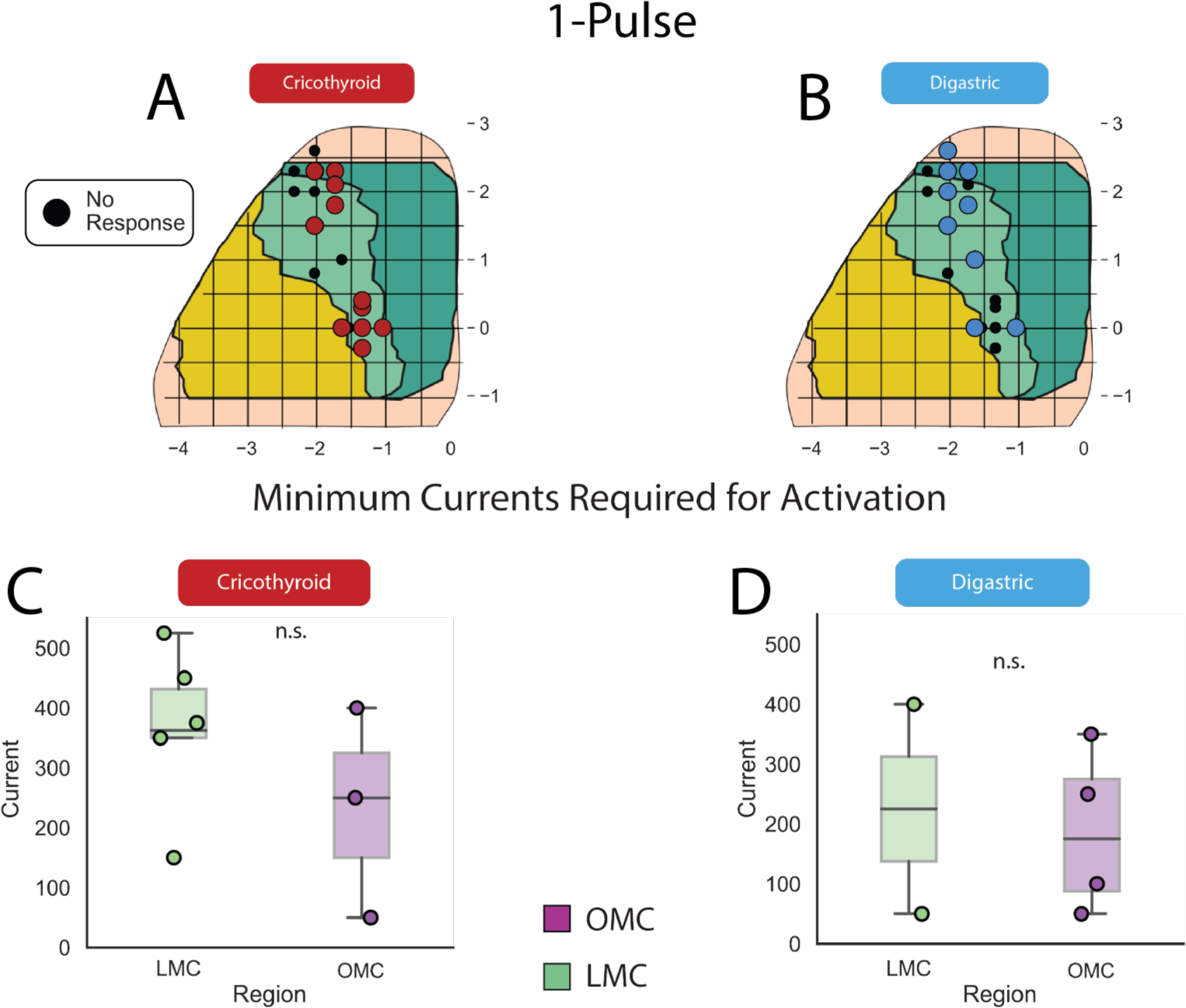
Single-Pulse Stimulated Sites and Minimum Response Currents. (A and B) Coordinates from all mice that were stimulated with single-pulse ICMS used to make kernel density estimates. Filled circles represent sites that were responsive for CT (red; A) and DG (blue; B). (C and D) Minimum currents across tested mice that resulted in a response from each muscle in each region. There was no significant different in the minimum current required for activation of each muscle (Mann-Whitney, p>0.05).

**Supplementary Figure 4.**
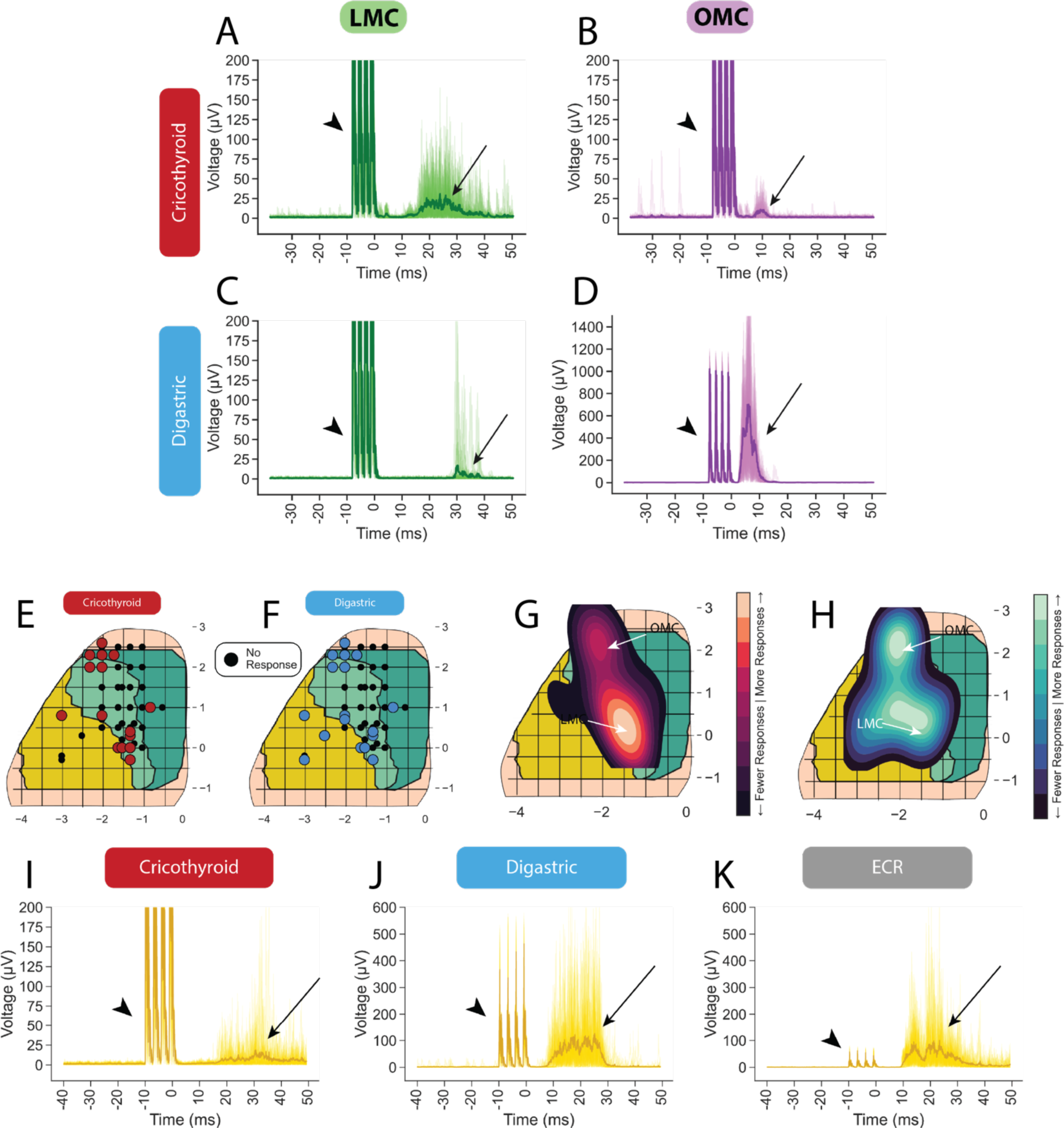
Responses of muscles from four-pulse stimulations. (A and B) Example traces and mean response of the CT muscle to a single round of four-pulse stimulations in LMC (A) and OMC (B). (C and D) Example traces and mean response of the DG muscle to a single round of four-pulse stimulations in LMC (C) and OMC (D). (E and F) Coordinates from all mice that were stimulated with four-pulse ICMS used to make kernel density estimates. Filled circles represent sites that were responsive for CT (red; E) and DG (blue; F). (G and H) Kernel density estimate weighted by percent of times the CT (G) and DG (H) muscle were responsive. (I-K) Example traces and mean response to stimulation at a coordinate in M1 (1.6L,1.6A) that activated the CT (I), DG (J), and ECR (K) muscles simultaneously.

**Supplementary Figure 5.**
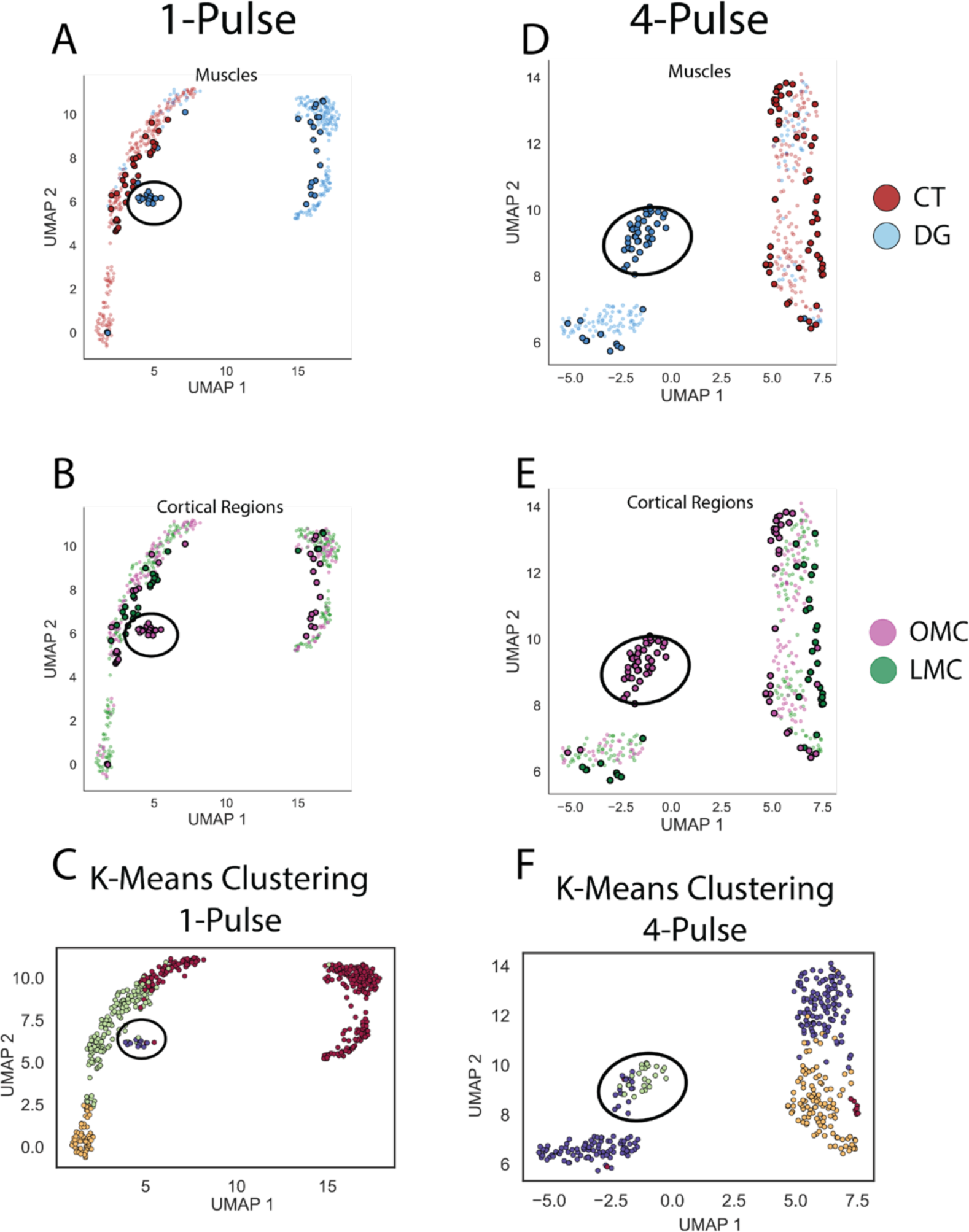
Low dimensional UMAP representations of StTAs. (A and B) UMAP projection of EMG StTA data. (A) is color coded by muscle and (B) is color coded by cortical region. Bolded points are responsive stimulations, pale points are non-responsive stimulations. (C) K-means clustering of the StTA data used to produce the UMAP plots (A and B). Points are color-coded by k-means cluster identity. Circled points represent the cluster identified by K-Means that represents the majority of DG responses from OMC in both single-pulse and four-pulse stimulations. (D-F) Similar representation of data as A-C from four-pulse stimulations. Information about muscle is more easily segregated than information about location of cortical stimulation.

**Supplementary Figure 6.**
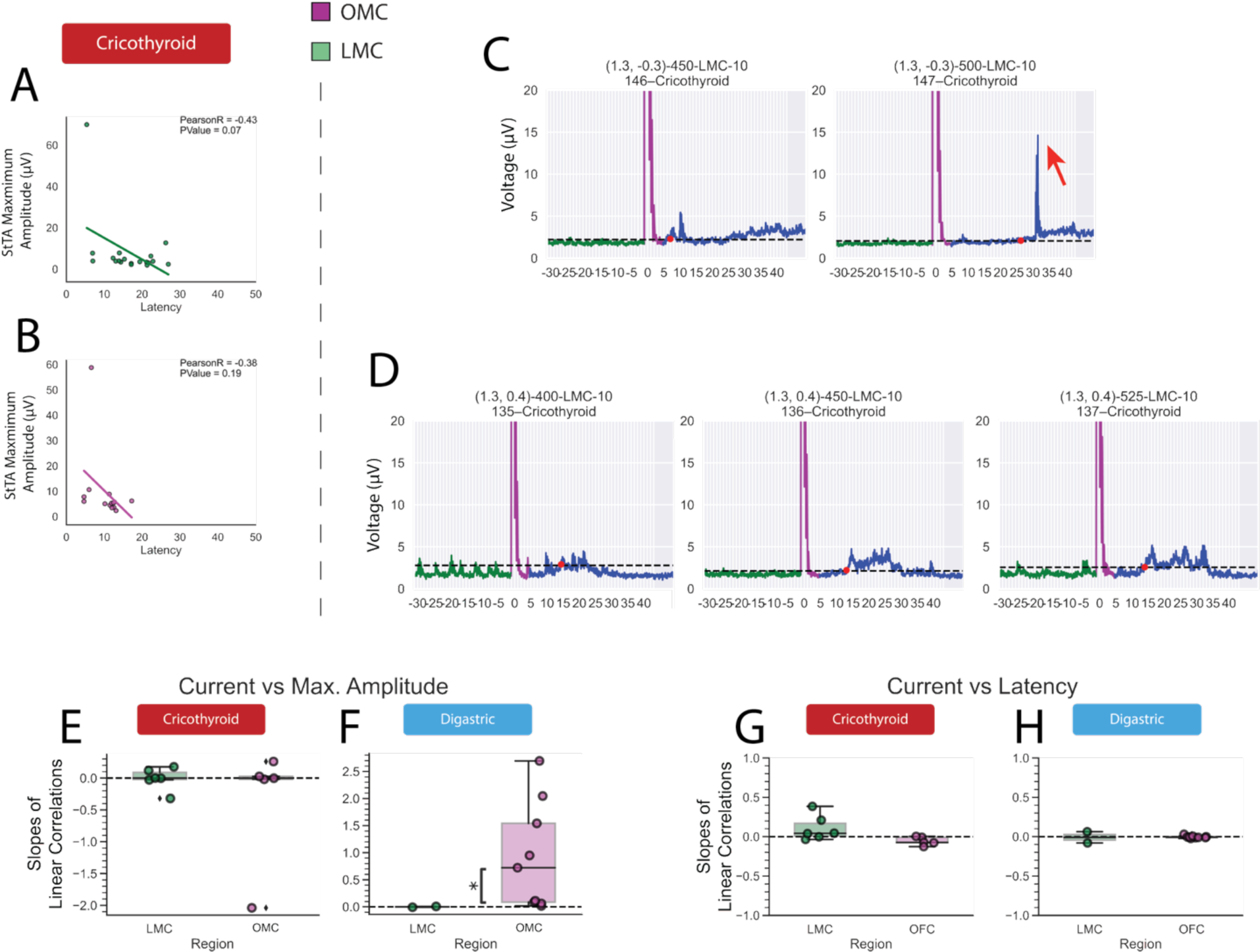
Relationship between latency, amplitude, and current in StTAs. (A and B) Relationship between latency and maximum amplitude for the CT muscle from stimulation in LMC (A) and OMC (B) without outliers removed (see Figure 3). (C) StTAs of a normal response on the left and the response with a recording artifact on the right that was removed as an outlier (red arrow). (D) Example set of StTAs from one coordinate at different currents without outlying measurements. StTA amplitude appears to subtly increase with current. (E and F) Slopes of correlations between current and maximum amplitude for the CT (E) and DG (F) muscles. In the CT responses, slopes were not significantly different from zero (Wilcoxon, p>0.05). In the DG responses, slopes in LMC were not significantly different from zero (Wilcoxon, p>0.05), and responses from OMC tended to be positive (Wilcoxon, p=0.0039). (G and H) Slopes of correlations between current and latency for the CT (G) and DG (H) muscles. In both the CT and DG responses, slopes were not significantly different from zero in either region (Wilcoxon, p>0.05).

**Supplementary Figure 7.**
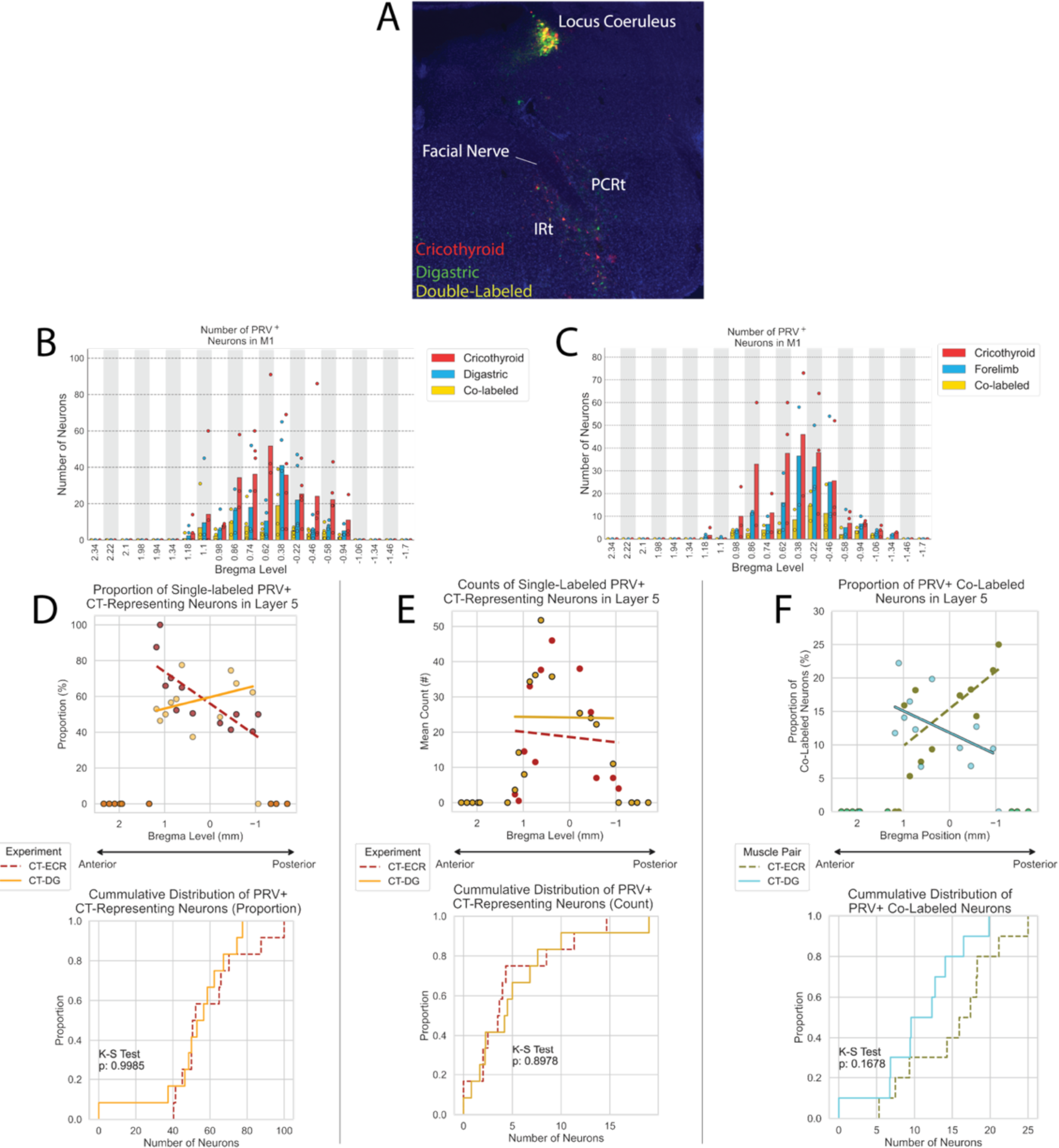
Neuron count from double-PRV injections. (A) Example of double-labeled neurons in locus coeruleus and the intermediate reticular formation (IRt). (B) Raw layer 5 neuron counts for each mouse with PRV injected into different muscles from Figure 4 C. (C) Raw neuron count for each mouse injected with PRV in CT and in ECR from Figure 5C. (D) Top: Proportion of single-labeled CT-representing neurons in each experiment. Single-labeled CT-representing neurons in the CT-FL experiment had a significant correlation with their anterior-posterior position (Pearson, p = 0.0026). Bottom: Cumulative distribution of the shared range of single-labeled CT-representing neurons. (E) Top: Mean number of single-labeled CT-representing neurons in each experiment. There was no significant correlation with position in either experiment. Bottom: Cumulative distribution of the shared range for the number of single-labeled CT-representing neurons. (F) Top: Proportion of double-labeled neurons (CT-FL or CT-DG) in each experiment. Double labeled CT-FL neurons had a significant correlation with their anterior-posterior position (Pearson, p = 0.026). Bottom: Cumulative distribution of the proportion of double-labeled neurons.

**Supplementary Figure 8.**
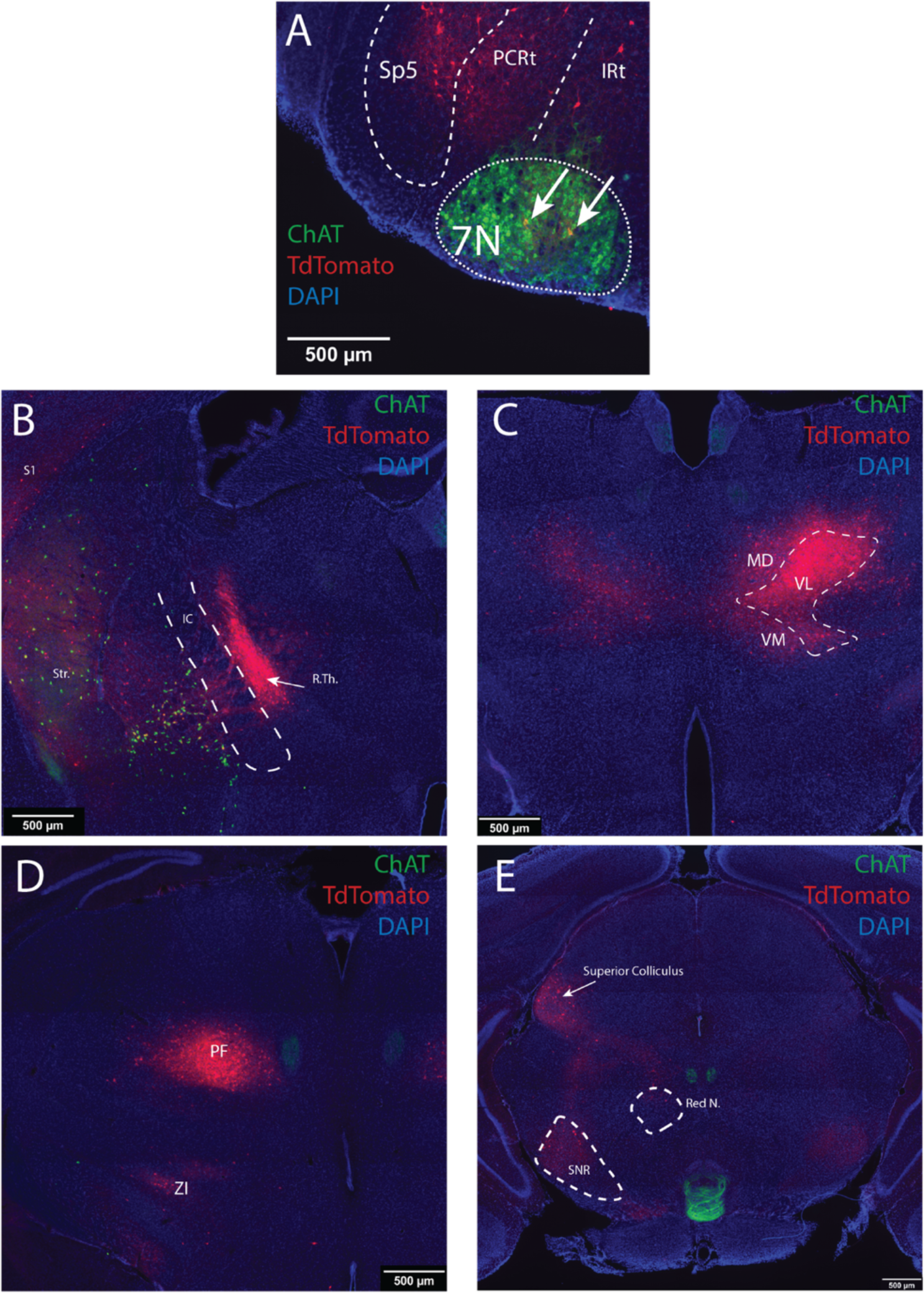
Other subcortical targets of OMC labeled post-transsynaptically with AAV1. (A) Example of post-transsynaptically labeled neurons (yellow and arrows) in the facial nucleus (7N). Verified with ChAT double-label (green). (B-D) Example image of post-transsynaptic label in different thalamic nuclei. Other striatal, S1, and zona inserta targets visible. (E) Example image of post-transsynaptic label to midbrain targets in the superior colliculus, substantia nigra (SNR), and the red nucleus. Anatomical abbreviations: Sp5, spinal trigeminal sensory nucleus; PCRt, parvicellular reticular formation; IRt, intermediate reticular formation; 7N, facial motor nucleus; S1, primary somatosensory cortex; Str., striatum; IC, internal capsule; R.Th., reticular thalamic nucleus; MD, mediodorsal thalamic nucleus; VL, ventrolateral thalamic nucleus; VM, ventromedial thalamic nucleus; PF, parafascicular nucleus; ZI, zona inserta; SNR, substantia nigra.

**Supplementary Figure 9.**
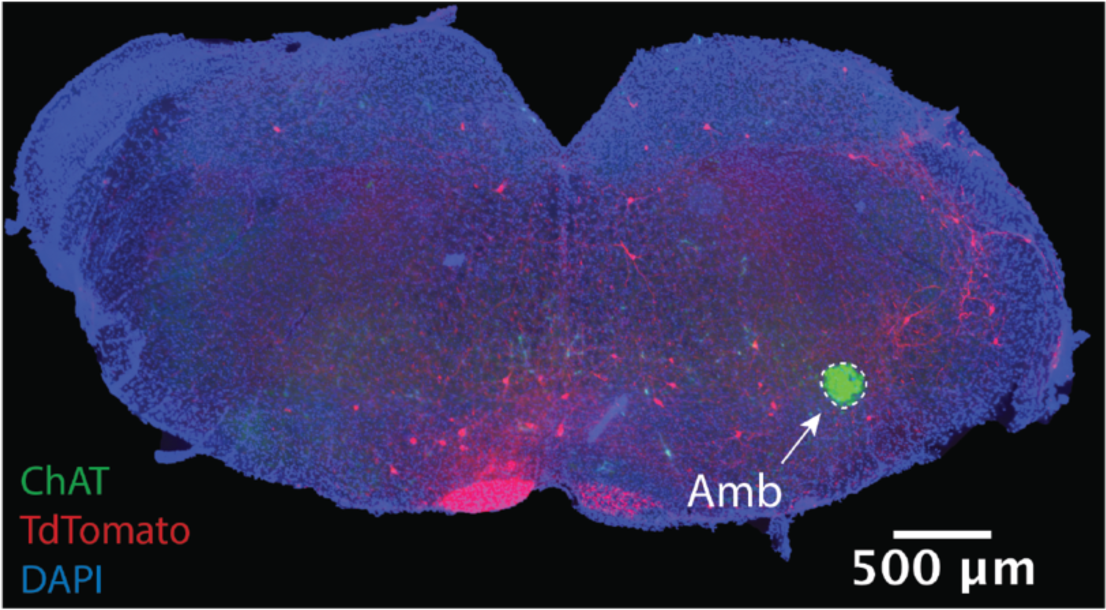
Lack of AAV1 post-transynaptic label from injection into LMC. Brainstem section at the level of Amb showing that AAV1 injections into LMC did not result in post-synaptically labeled targets in Amb despite the known direct projection.

**Supplementary Figure 10.**
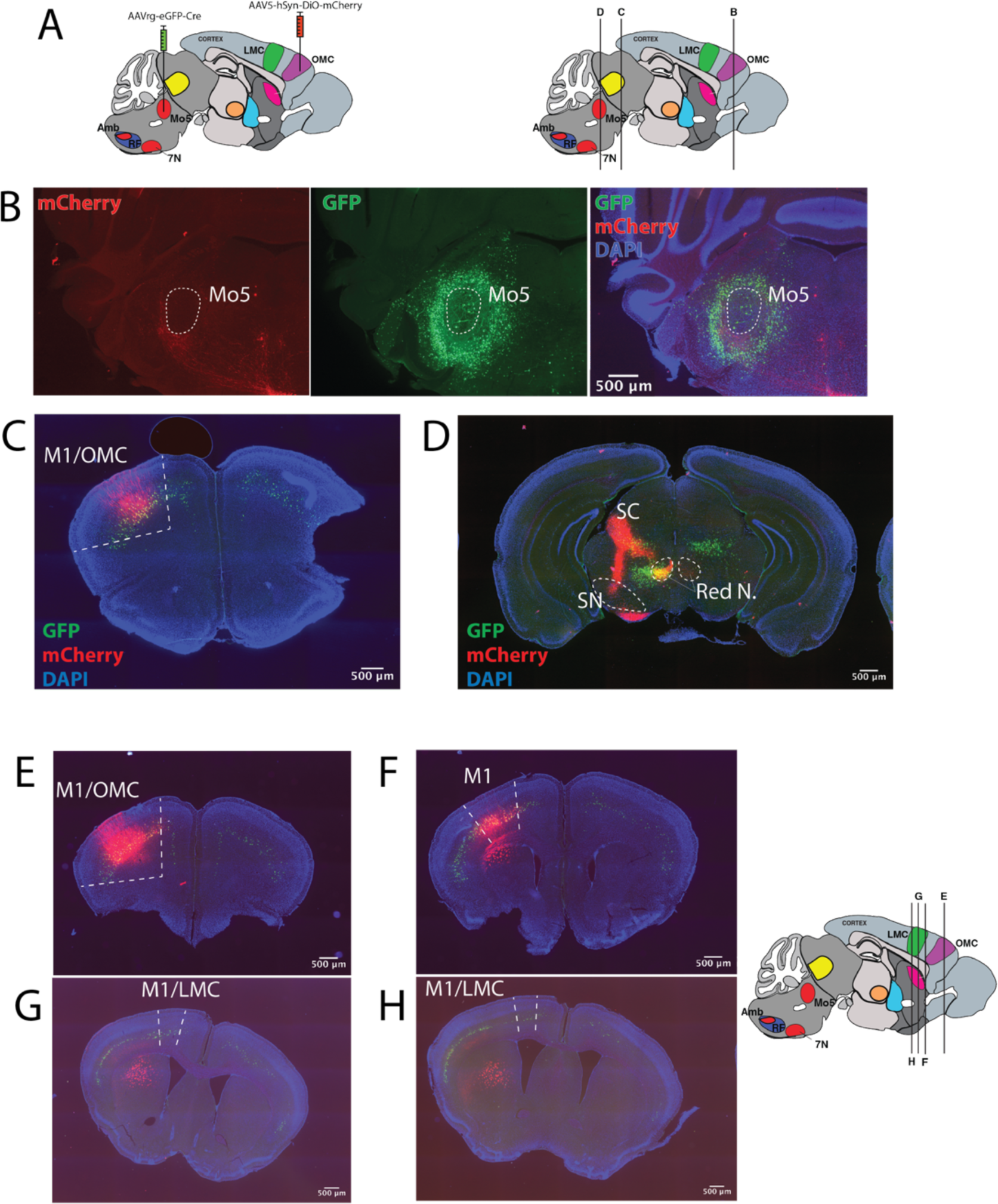
Population of direct-projecting OMC neurons make many distal collateral connections. (A) Overview of viral strategy (B) Example image of terminal fibers originating in OMC (red; Cre-mediated mCherry) and neurons projecting to Mo5 (green; eGFP-Cre). (C) Mo5-projecting neurons in OMC whose terminals are visible in panel (B). (D) Example of collateral fibers in midbrain superior collicular targets which originate from the same population of OMC neurons that project to Mo5. Some convergence of non-cortical Mo5-projecting neurons overlap with OMC collaterals. (E-H) Intracortical projection targets of Mo5-projecting OMC neurons to S1 (red). Note lack of fibers in LMC region. Some retrogradely labeled neurons in S1 from Mo5 injection (green).

**Supplementary Figure 11.**
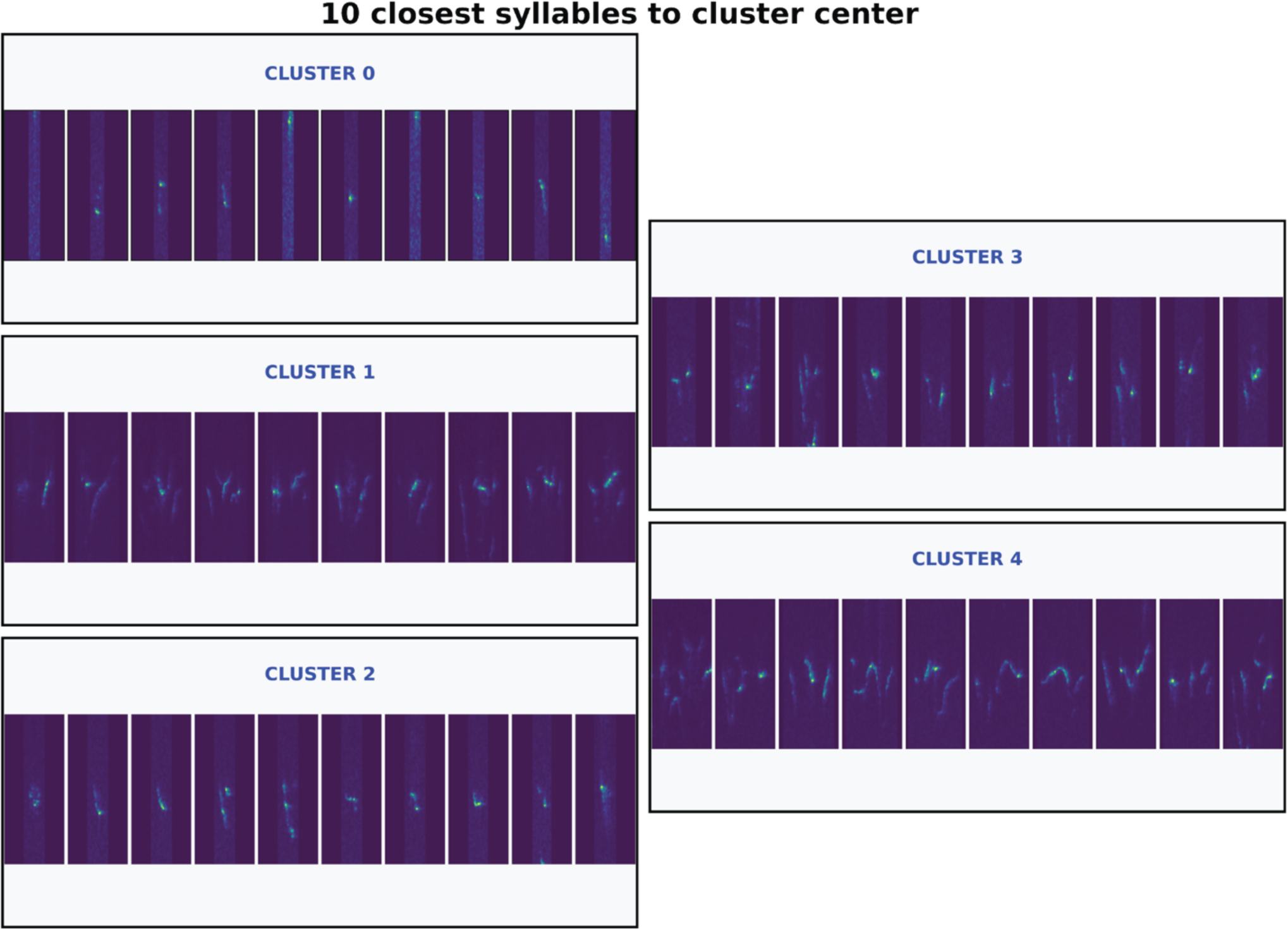
Unsupervised Clusters used in Syntax Analysis. Syllable clusters determined by unsupervised clustering in AMVOC. Each set of images contains the ten syllables closes to the center of the clustering distribution.

